# Interleukin-1 receptor antagonist is a conserved factor for exacerbating tuberculosis susceptibility

**DOI:** 10.1101/2023.10.27.564420

**Authors:** Ophelia V. Lee, Enish Pathak, Sujhey Hernández-Bazán, Daisy X. Ji, Bruce A. Rosa, David L. Jaye, Sara Suliman, Makedonka Mitreva, Cem Gabay, Russell E. Vance, Dmitri I. Kotov

**Affiliations:** Divison of Immunology and Molecular Medicine, University of California, Berkeley, Berkeley, CA, 94720, USA; Division of Infectious Diseases, Department of Medicine, Washington University School of Medicine, St. Louis, MO, 63110, USA; Department of Microbiology, New York University Grossman School of Medicine, New York, NY, 10016, USA; Department of Pathology and Laboratory Medicine, Emory University, Atlanta, GA, 30322, USA; Division of Experimental Medicine, Department of Medicine, University of California, San Francisco, San Francisco, CA, 94115, USA; Division of Rheumatology, Department of Medicine, University Hospital of Geneva, Geneva, Switzerland; Divison of Immunology and Molecular Medicine, Howard Hughes Medical Institute, University of California, Berkeley, Berkeley, CA, 94720, USA; Andrew M. and Jane M. Bursky Center for Human Immunology and Immunotherapy Programs, Washington University School of Medicine, St. Louis, MO, 63110, USA

**Author notes:** These authors contributed equally (Ophelia V. Lee, Enish Pathak). Lead contact: Requests for further information and resources should be directed to and will be fulfilled by the lead contact, Dmitri I. Kotov.

**Keywords:** Innate immunity, tuberculosis, IL-1 receptor antagonist, neutrophil, macrophage, osteopontin, TNF, Spp1

## Abstract

*Mycobacterium tuberculosis* (*Mtb*) causes 1.25 million deaths a year; however, tuberculosis (TB) pathogenesis remains poorly understood. Here we find that gene signatures from three different *Mtb*-susceptible mouse models predict active TB disease in humans significantly better than a signature from resistant C57BL/6 (B6) mice. Conserved among susceptible mice, non-human primates, and humans, but largely absent from B6 mice, was *Mtb*-induced *Spp1*^+^ macrophage differentiation. *Spp1*^+^ macrophages expressed high levels of immunosuppressive molecules including IL-1 receptor antagonist (IL-1Ra). Here we report that enhancement of IL-1 signaling via deletion of IL-1Ra promoted bacterial control across three susceptible mouse models. We found that IL-1 signaling promotes pulmonary control of *Mtb* infection through amplifying TNF production by uninfected bystander cells. Our results indicate that myeloid cell expression of immunosuppressive molecules, in particular IL-1 receptor antagonist, is a conserved mechanism limiting *Mtb* control in mice, non-human primates, and humans.

## Introduction

*Mycobacterium tuberculosis* (*Mtb*) causes ∼10 million new cases of active tuberculosis (TB) disease and 1.25 million deaths per year, and is the leading cause of mortality from a bacterial pathogen^1^. Unfortunately, TB treatment consists of at least a 4-6 month regimen of antibiotics, and the only approved vaccine has little to no efficacy in adults^2^. A greater understanding of the infection is essential, but this depends on tractable animal models that appropriately mimic *Mtb* pathogenesis in humans. While mouse models have led to key discoveries such as the critical role of IFN-γ^3,4^ and TNF^5^ in *Mtb* control, mouse models have also been criticized for not recapitulating key aspects of human disease, such as highly structured granuloma formation^6,7^. Further hallmarks of active TB disease in humans include expression of type I interferon (IFN) stimulated genes and a heightened neutrophil response^8,9^. The standard, resistant C57BL/6 (B6) mouse model only weakly exhibits these hallmarks^10–13^.

We recently described *Sp140*^-/-^ mice as an *Mtb*-susceptible mouse model that mimics the exacerbated type I IFN signature and heightened neutrophil response seen in humans^14,15^. *Sp140* encodes a transcriptional repressor that limits expression of the canonical type I IFN gene *Ifnb1*^16^. Loss of *Sp140* accounts for the excessive type I IFN response and *Mtb*-susceptibility of the previously described *Sst1*^S^ (“Kramink”) mice^17–19^. Deletion of the gene encoding the type I IFN receptor (*Ifnar1*) rescues the hypersusceptibility of both *Sst1*^S^ and *Sp140*^-/-^ mice, implying a causal role of type I IFNs in *Mtb* pathogenesis in these models^14,15^. Other *Mtb* susceptible mouse models have been described, such as *Nos2*^-/-^ and *Acod1*^-/-^ mice^20,21^, though it remains unclear whether disease in these models is also driven by type I IFN signaling. Nevertheless, we hypothesized that there may be conserved features underlying the susceptibility of diverse mouse models, and that these features may be preserved across susceptible mice, non-human primates, and humans.

Neutrophilic inflammation is a major feature associated with TB disease across *Sp140*^-/-^, *Nos2*^-/-^, *Acod1*^-/-^ and other susceptible mouse models^13,15,21–24^, as well as during non-human primate^25,26^ and human TB^27,28^. Depletion of neutrophils enhances bacterial control in diverse *Mtb*-susceptible mouse models, including *Sp140*^-/-^, *Nos2*^-/-^, and *Acod1*^-/-^ mice^13,15,21^. However, the precise role of neutrophils in driving bacterial replication is unclear. One widely held view is that neutrophil necrosis and inflammation creates a replicative niche for the bacteria, including possibly within neutrophils themselves^29,30^. The production of neutrophil extracellular traps (NETs) is also speculated to promote bacterial replication, perhaps by induction of type I IFNs^15,22,31^. In states of chronic inflammation, neutrophils can also play immunosuppressive roles^32^, though it remains uncertain if this accounts for neutrophil-driven exacerbation of TB disease_13,15,21_.

Signaling induced by interleukin-1 (IL-1, comprising IL-1α and IL-1β) potently recruits neutrophils to sites of inflammation, and IL-1 is expressed at high levels in *Mtb*-susceptible mice^17,33^ and humans^34^. Excessive IL-1 has been proposed to enhance bacterial replication^35^, particularly in *Nos2*^-/-^ mice^13,33^, though the underlying mechanisms remain uncertain. A pro-bacterial function of IL-1 is difficult to reconcile with its clear anti-bacterial functions, as mice deficient in IL-1 signaling are highly susceptible to *Mtb* infection^23,36–40^. Importantly, the levels of IL-1 do not necessarily correlate with its signaling capacity. This is because IL-1 signaling is repressed by a decoy receptor (IL-1R2) and a soluble receptor antagonist (IL-1Ra, encoded by the *Il1rn* gene). Although susceptible *Sst1*^S^ mice express high levels of IL-1α and IL-1β, we previously reported that deficiency in *Il1rn* (resulting in enhanced IL-1 signaling) markedly promotes bacterial control in these mice^17^. These results imply that *Mtb*-susceptibility can arise from a lack of IL-1 signaling, rather than excessive IL-1 signaling, even when IL-1 levels are high. However, disease in *Sst1*^S^ mice may be a unique case since it is driven by an exacerbated type I IFN response^17^, and *Il1rn* is an interferon-stimulated gene. Thus, it remains unclear whether impairment of IL-1 signaling by IL-1Ra plays a conserved role in humans or other *Mtb*-susceptible mouse models, especially *Nos2*^-/-^ and *Acod1*^-/-^ mice, in which *Mtb* infection is characterized by high levels of IL-1.

Here, in an effort to identify conserved mechanisms underlying *Mtb* susceptibility, we used single cell RNA sequencing (scRNA-seq) to profile and characterize three different *Mtb*-susceptible mouse strains (*Sp140*^-/-^, *Nos2*^-/-^, *Acod1*^-/-^ mice). Unlike *Sp140*^-/-^ mice, we found that *Nos2*^-/-^ and *Acod1*^-/-^ mice did not exhibit a strong type I IFN signature and were not rescued by *Ifnar1* deficiency. Despite this, a common feature of *Mtb*-induced genes across all three strains was myeloid cell expression of immunosuppressive genes. In support of the importance of shared features of the *Mtb*-susceptible strains, gene signatures from each of these strains predicted human TB progression better than a signature obtained from resistant B6 mice. Additionally, macrophage differentiation into an *Spp1*^+^ state was conserved across the susceptible mouse strains, *Mtb*-infected non-human primates, and *Mtb*-infected humans.

Mouse, non-human primate, and human *Spp1*^+^ macrophages expressed immunosuppressive molecules, including *Il1rn*. Given the importance of IL-1 signaling for *Mtb* control, and the strong expression of IL-1Ra in myeloid cells, we focused on the role of IL-1Ra in *Mtb* susceptibility.

Global deletion as well as myeloid-specific deletion of IL-1Ra rescued bacterial control in *Sp140*^-/-^ mice, demonstrating a major role for macrophage expression of this immunosuppressive molecule in driving *Mtb* disease. This contribution was shared across the susceptible mouse models, as global IL-1Ra deletion also enhanced bacterial control in *Nos2*^-/-^ and *Acod1*^-/-^ mice while neutrophil-specific IL-1Ra deletion did not, in line with macrophages being the dominant source of IL-1Ra during *Mtb* infection. Although *IL1RN* has been included in multiple human TB gene signatures^8,41–43^, we find it alone was sufficient to strongly predict human TB and was induced ∼200 days prior to TB diagnosis. Together, these data suggest that immunosuppression by myeloid cells is a conserved feature of *Mtb* susceptibility across mice and humans, with some immunosuppressive pathways, such as IL-1 inhibition via IL-1Ra, being a shared mechanism aggravating TB.

## Results

### Mtb-susceptible mice more accurately model human infection than B6 mice

Human *Mtb* infection presents as a highly diverse spectrum of diseased and non-diseased states. The best characterized hallmarks of active disease include induction of type I interferon (IFN) genes and a myeloid cell inflammatory response, especially by neutrophils^8^. We previously demonstrated that *Sp140*^-/-^ mice exhibit both hallmarks, and that these hallmarks drive susceptibility in these mice^14,15^. Neutrophil trafficking to the lungs also promotes *Mtb* susceptibility in other mouse models^22,44–46^, including *Nos2*^-/-^ and *Acod1*^-/-^ mice^13,21^. We sought to explore the use of different susceptible mouse models as a way to mimic the diversity of human TB disease. To start, we characterized the myeloid cell response in *Sp140*^-/-^, *Nos2*^-/-^, and *Acod1*^-/-^ mice. We focused on ∼4 weeks post-infection since it was a relatively early timepoint by which clear differences were reported between *Mtb* resistant B6 mice and susceptible strains^13–15,21^. Although there are also major differences between strains at later timepoints, our rationale was to focus on earlier events to try to identify more proximal innate immune determinants of susceptibility.

In line with previous reports, we observed a ∼5-fold increase in lung neutrophils and a ∼2-4-fold increase in lung monocytes and interstitial macrophages (IMs) in *Mtb*-susceptible mice 25 days after *Mtb* infection in each of the three mouse models (**Extended Data Fig. 1A**). The increase in lung neutrophils was also observed as a proportional increase in neutrophils among total lung cells in all three *Mtb*-susceptible models (**Extended Data Fig. 1B**). We sought to explore myeloid function in promoting *Mtb* susceptibility by first examining myeloid cell trafficking in *Mtb*-infected lungs from B6 and *Sp140*^-/-^ mice (**Extended Data Fig. 1C**). Diseased tissue within the lung was defined by the presence of *Mtb* and high cellularity. These features were largely absent from healthy portions of the lung. Diseased areas of the lungs had more macrophages (SIRPα^+^ cells) and neutrophils (Ly6G^+^ cells) than the healthy tissue of the same lungs (**Extended Data Fig. 1D**). To confirm that this enrichment was not unique to *Sp140*^-/-^ mice, we also compared macrophage and neutrophil trafficking in B6, *Nos2*^-/-^, and *Acod1*^-/-^ mice (**Extended Data Fig. 1E**). Similar to *Sp140*^-/-^ mice, neutrophils were significantly enriched, and macrophages trended toward greater numbers in diseased relative to healthy tissue in all tested genotypes (**Extended Data Fig. 1F**). Interestingly, while similar numbers of macrophages and neutrophils were recruited to the diseased portions of susceptible mouse lungs, macrophages were significantly closer to *Mtb* in B6, *Sp140*^-/-^, *Nos2*^-/-^, and *Acod1*^-/-^ mice, as compared to neutrophils, which were more spatially segregated from *Mtb* (**Extended Data Fig. 1G**). These data indicate that myeloid cells are more abundant in *Mtb*-infected lungs and preferentially traffic to sites of disease within those lungs, with macrophages residing closer to *Mtb* within the infected lung tissue.

As the heightened myeloid response was conserved across the *Mtb*-susceptible mouse models, we sought to test whether *Nos2*^-/-^ and *Acod1*^-/-^ mice share other attributes of the susceptibility of *Sp140*^-/-^ mice, including type I IFN-driven disease. Myeloid cells are the primary reservoir of *Mtb*, so we used our previously described single cell RNA-sequencing (scRNA-seq) workflow^15^ to profile myeloid cells in *Mtb*-infected and naïve *Nos2*^-/-^ and *Acod1*^-/-^ mice. Infection with a fluorescent (Wasabi^+^) *Mtb* strain allowed us to distinguish infected and bystander cells in infected mice. The resulting datasets were integrated with our previously published B6 and *Sp140*^-/-^ myeloid cell datasets^15^ using the R package Seurat^47^ (**Fig. 1A**). B6 mice were used as a control rather than *Sp140*^-/-^ *Ifnar1*^-/-^ mice because we previously demonstrated there is no difference in bacterial burden between these two genotypes 25 days post-infection (the timepoint of the scRNA-seq experiment)^14^. As these are CITE-seq^48^ datasets, mRNA and protein expression of lineage and activation markers were implemented for the manual annotation of cell clusters (**Extended Data Fig. 2A, 2B**). Comparing the myeloid cells from naïve lungs of B6, *Sp140*^-/-^, *Nos2*^-/-^, and *Acod1*^-/-^ mice revealed very few differentially expressed genes between the mouse lines, suggesting that each genotype had a similar inflammatory state prior to infection (**Extended Data Fig. 2C, 2D**). Therefore, differences between genotypes in *Mtb*-infected mice were likely induced in response to the infection rather than the result of pre-existing inflammation. Analysis of the myeloid cell clusters within the integrated dataset showed that the dominant myeloid cell populations (monocytes, macrophages, and neutrophils) are present in each genotype.

**Figure 1.**
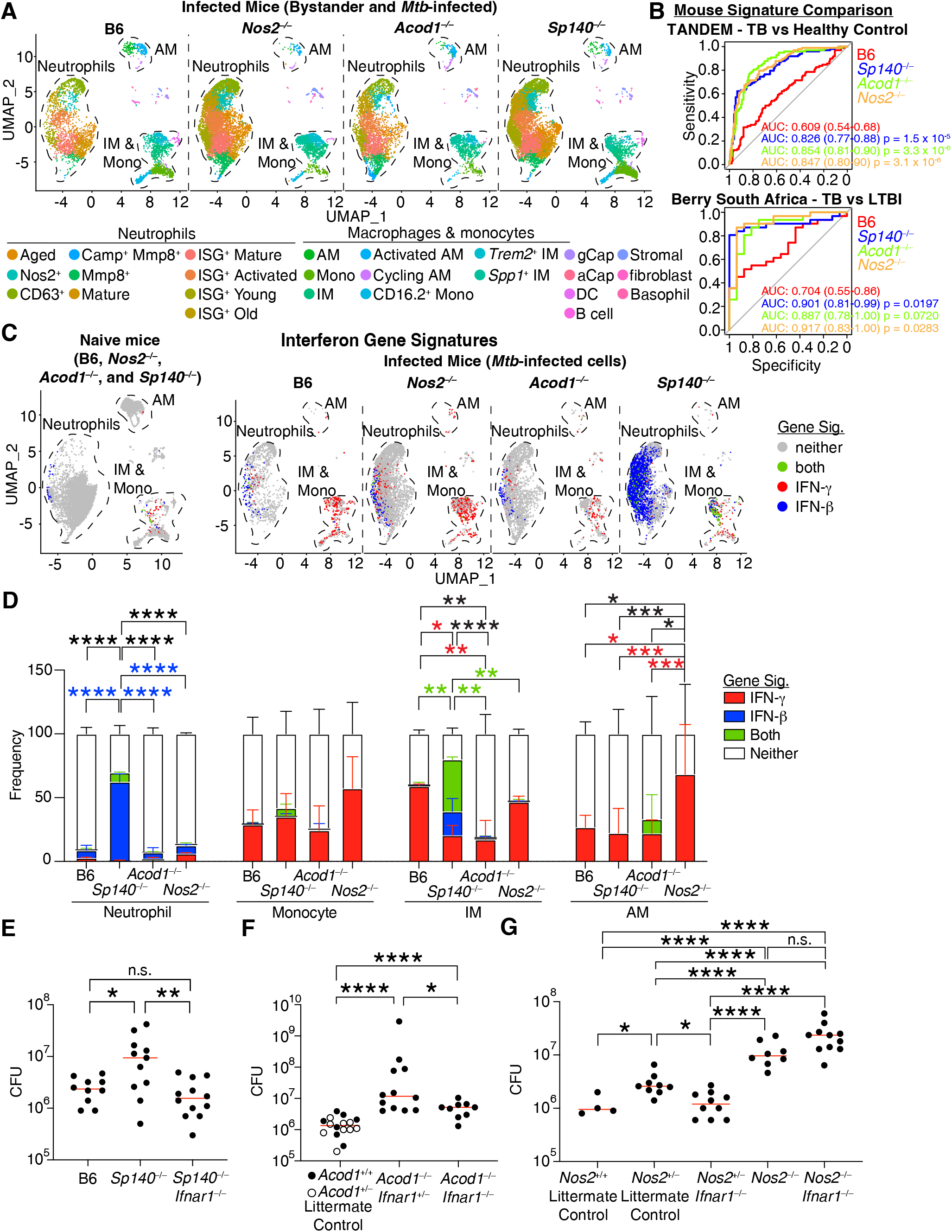
*Nos2*-and *Acod1*-deficient mice mimic human *Mtb* progression but do not exhibit type I interferon-driven susceptibility. (**A**) Visualizing lung myeloid cell populations in a scRNA-seq dataset of cells from *Mtb*-infected B6, *Nos2*^-/-^, *Acod1*^-/-^, and *Sp140*^-/-^ mice (n = 3 per genotype). (**B**) ROC curves of the ability of genes induced by *Mtb* infection in B6, *Nos2*^-/-^, *Acod1*^-/-^, or *Sp140*^-/-^ mice to stratify human active *Mtb* patients (n = 151) and healthy controls (n = 98) in the TANDEM cohort (analysis of GSE114192) as well as stratifying human active *Mtb* patients (n = 16) and LTBI individuals (n = 31) in the Berry South Africa cohort (analysis of GSE107992). Parenthetical numbers indicate the 95% confidence interval, and p-values are for comparisons between the given susceptible mouse signature and the B6 signature. (**C**) UMAP plot and (**D**) quantification of myeloid cells undergoing type I interferon and/or IFN-γ signaling in the lungs of naïve or *Mtb*-infected B6, *Nos2*^-/-^, *Acod1*^-/-^, or *Sp140*^-/-^ mice (n = 3 per genotype and infection status). (**E**) Lung bacterial burden in B6 (n = 10), *Sp140*^-/-^ (n = 11), and *Sp140*^-/-^ *Ifnar1*^-/-^ (n = 12) mice infected with *Mtb*. (**F**) Lung bacterial burden in littermate control (*Acod1*^+/+^ and *Acod1*^+/-^ mice combined due to their equal *Mtb* burden, n = 16), *Acod1*^-/-^ *Ifnar1*^+/-^ (n = 12), and *Acod1*^-/-^ *Ifnar1*^-/-^ (n = 9) mice infected with *Mtb*. (**G**) Lung bacterial burden in *Nos2*^+/+^ littermate control (n = 4), *Nos2*^+/-^ littermate control (n = 9), *Nos2*^+/-^ *Ifnar1*^-/-^ (n = 10), *Nos2*^-/-^ (n = 8), and *Nos2*^-/-^ *Ifnar1*^-/-^ (n = 11) mice infected with *Mtb*. Lungs were analyzed 25-27 days after *Mtb*-Wasabi infection. The error bars in (D) represent the standard deviation. The bars in (E), (F), and (G) represent the median. Pooled data from two independent experiments are shown in (E), (F), and (G). Statistical significance in (D), (E), (F), and (G) by one-way ANOVA with Tukey’s multiple comparison test and in (B) by Bootstrap test. *p < 0.05, **p < 0.01, ***p < 0.001, ****p < 0.0001.

Given that the increase in neutrophils exhibited by *Mtb*-susceptible mouse models is a hallmark of human *Mtb* disease progression^27^, we hypothesized that susceptible mouse models might mimic additional conserved hallmarks of active human disease, in which case they could be a useful tool to study *Mtb* pathogenesis. To test this hypothesis, we identified lung myeloid gene signatures consisting of genes induced at least 8-fold in *Mtb*-infected mice relative to naïve lungs (**Extended Data Fig. 3A**). The ability of these genes to discriminate *Mtb* cases from healthy controls in the TANDEM cohort^49^ was quantified by calculating the area under the curve (AUC) for receiver operating characteristic (ROC) curves, with values closer to 1 indicating better classifiers of human disease (**Fig. 1B**; p-values compare performance of *Mtb*-susceptible mouse signatures to the B6 signature). While there was no major difference in the ability of the signatures derived from the three susceptible mouse models to discern human disease, all the susceptible models considerably outperformed the signature derived from B6 mice, suggesting the susceptible mouse models better recapitulate characteristics of active human disease than B6 mice. We also tested the ability of these signatures to discriminate between active TB and latent TB-infected (LTBI) individuals in the Berry South Africa cohort^50^, and similarly observed that the three susceptible mouse model signatures out-performed the B6 mouse signature, in line with the hypothesis that susceptible mouse models better recapitulate active human disease (**Fig. 1B**). We corroborated this finding by generating new signatures for each genotype through machine learning feature selection using a Random Forest algorithm with the TANDEM cohort as the training dataset and the Berry South Africa cohort as the test dataset. The machine learning-generated signatures for the susceptible mice also significantly outperformed the new B6 mouse signature in discerning active TB from LTBI individuals, supporting the idea that susceptible mice better recapitulating features of human disease (**Extended Data Fig. 3B**). The goal of this analysis was to validate the utility of using these mouse models to understand TB pathogenesis in humans rather than to identify potential biomarkers of human TB disease.

However, we also observed that the mouse signatures discriminated active TB infection from lung cancer and pneumonia in the Bloom et al. cohort^51^, suggesting the similarity in immune responses between mice and humans captured by these gene signatures does not solely reflect general lung inflammation but captures TB-specific pathways (**Extended Data Fig. 3C**).

Next, we sought to better understand why the susceptible models were all able to discriminate between *Mtb* cases and healthy controls by assessing whether the type I IFN response present in humans^8^ and *Sp140*^-/-^ mice^14,15^ was also present in *Nos2*^-/-^ and *Acod1*^-/-^ animals. Using our previously published gene signatures that distinguish between type I IFN and IFN-γ signaling^15^, we classified cells as responding to type I IFN and/or IFN-γ in our myeloid scRNA-seq dataset. As expected, very few cells from naïve lungs were positive for either the type I or II IFN signature (**Fig. 1C**). Conversely, at least some *Mtb*-infected cells from every genotype had strong IFN-γ signaling, especially in monocytes and macrophages (**Fig. 1C, 1D**). However, the type I IFN response was notable only in the *Sp140*^-/-^ mice, with strong type I IFN signaling in the neutrophils and interstitial macrophages (IMs) of these mice.

We experimentally tested whether type I IFN signaling plays a causative role in disease progression in each *Mtb*-susceptible mouse model by examining lung bacterial burden in *Ifnar1*-deficient *Sp140*^-/-^, *Nos2*^-/-^, and *Acod1*^-/-^ mice 25 days after *Mtb* infection. As reported previously, global *Ifnar1*-deficiency fully rescued the susceptibility of *Sp140*^-/-^ mice, phenocopying the *Mtb* restriction of B6 mice^14,15^; however, *Ifnar1*-deficiency only slightly reduced bacterial burdens in *Acod1*^-/-^ mice and even modestly but not statistically significantly increased bacterial burdens in *Nos2*^-/-^ mice (**Fig. 1E, 1F, 1G**). Together, these data suggest that while all three models recapitulate key aspects of human TB, type I IFN-driven susceptibility was largely unique to *Sp140*^-/-^ mice, indicating other mechanisms can also promote TB pathogenesis.

### Myeloid cell production of IL-1Ra is a conserved hallmark of Mtb susceptibility

As a strong type I IFN response was not shared among the three *Mtb*-susceptible mouse models, we analyzed our scRNA-seq to identify common features that may contribute to the elevated bacterial burdens present in these animals. We observed that *Mtb* infection resulted in activation of the IMs, driving them to adopt either a proinflammatory *Trem2*^+^ or an alternative *Spp1*^+^ activation state^52–54^ (**Fig. 2A, 2B**). Interestingly, *Spp1*^+^ macrophages were the dominant monocyte-derived macrophage population in the *Mtb*-susceptible *Sp140*^-/-^, *Nos2*^-/-^, and *Acod1*^-/-^ mice, while being largely absent in *Mtb*-restrictive B6 mice (**Fig. 2C, 2D**). The *Spp1*^+^ state was distinguished from the *Trem2*^+^ state by the elevated expression of immunosuppressive molecules in *Mtb*-harboring macrophages, including *Il1rn* (encoding IL-1Ra) which we previously demonstrated contributes to loss of bacterial control in *Mtb*-susceptible *Sst1*^S^ mice^17^ (**Fig. 2E**). Alongside neutrophils, which constitutively express *Il1rn*^55^, *Spp1*^+^ IMs were the dominant source of *Il1rn* among myeloid cells and *Spp1*^+^ IM *Il1rn* expression was elevated in all three susceptible strains relative to B6 mice (**Fig. 2F**). Reanalysis, including re-annotation, of the myeloid cells in a published scRNA-seq dataset of non-human primate *Mtb* granulomas^56^ indicated that *SPP1^+^* macrophages were also present in non-human primate *Mtb* granulomas (**Fig. 2G, Extended Data Fig. 4A**). These *SPP1^+^* macrophages expressed higher levels of immunosuppressive molecules such as *IDO1*, *CD274*, *IL18BP*, and *IL1RN* than *TREM2^+^* macrophages (**Fig. 2H**). Reanalysis and re-annotation of the myeloid portion of a scRNA-seq dataset of resected lungs from *Mtb*-infected humans^57^ also identified SPP1^+^ IMs with elevated expression of immunosuppressive molecules relative to TREM2^+^ IMs, including increased expression of *IDO1*, *TGFB1*, *IL10*, *IL27*, *CD274*, *IL1RN*, and the IL-1 decoy receptor *IL1R2* (**Fig. 2I, 2J**; **Extended Data Fig. 4B**). Additionally, *SPP1^+^* IMs were a dominant source of *IL1RN* among myeloid cells in the human lungs with the *IL1RN*-expressing *SPP1^+^* IMs present in metabolically active (FDG high signal) and metabolically inactive (FDG low signal) portions of *Mtb*-infected human lungs (**Fig. 2K**). Further examining the conserved nature of *Spp1*^+^ macrophages, we identified a core *Spp1*^+^ gene signature consisting of genes upregulated in *Spp1*^+^ relative to *Trem2*^+^ IMs in the three susceptible mouse models as well as in human *Mtb*-infected lungs (**Extended Data Fig. 4C**). This core signature includes *IL1RN,* highlighting the strong association between *Spp1*^+^ IM differentiation and *IL1RN* expression across species (**Extended Data Fig. 4D**). Moreover, we observed *IL1RN* is elevated in *Mtb*-harboring myeloid cells from infected mice as well as the PBMCs of *Mtb*-infected humans (**Extended Data Fig. 5A**, **5B**, **5C**).

**Figure 2.**
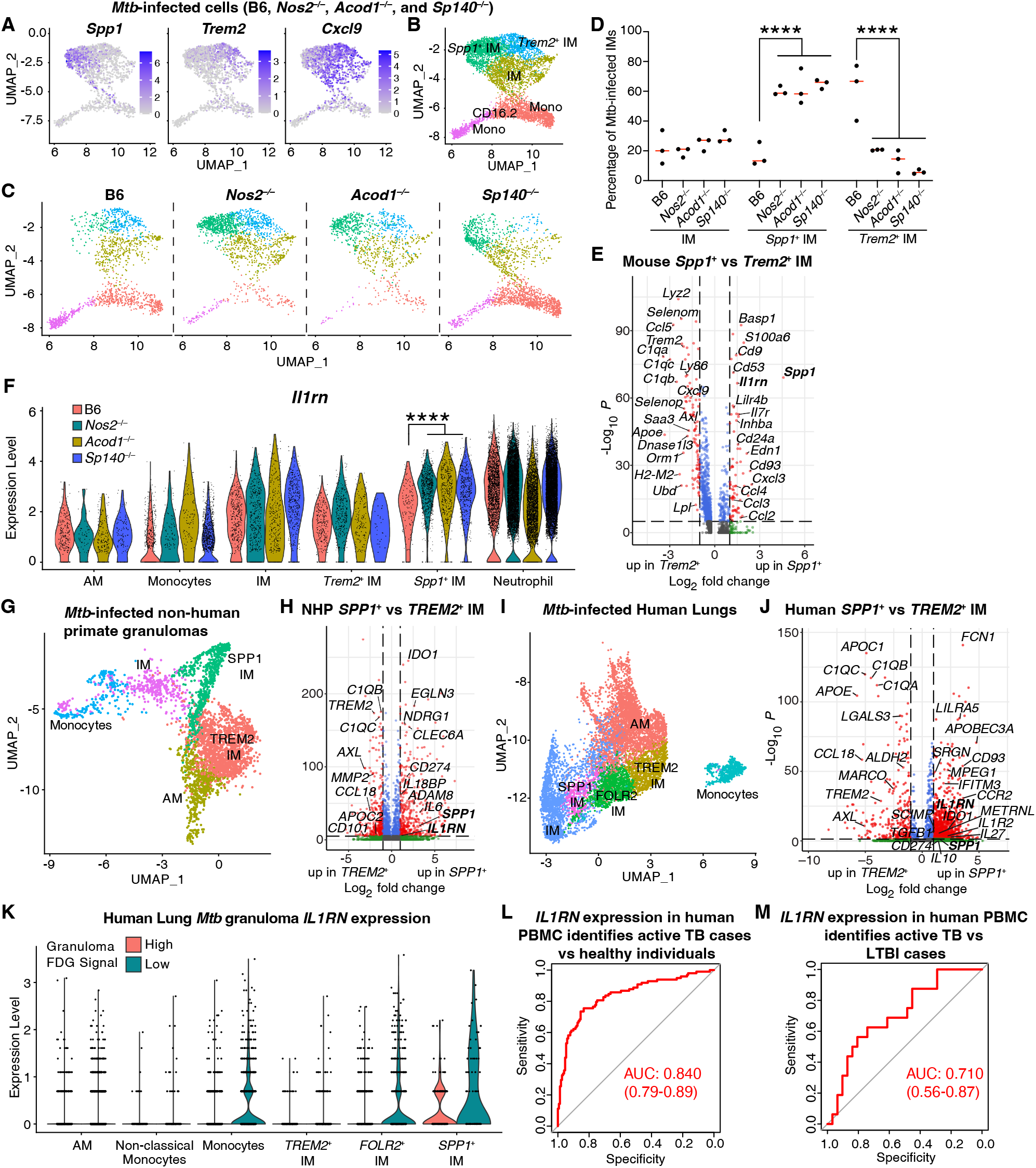
*Spp1*^+^ macrophage differentiation and their expression of immunosuppressive molecules, including IL-1 receptor antagonist, correlates with *Mtb* susceptibility in mice and is conserved between mice, *Mtb*-infected non-human primates, and *Mtb*-infected humans. (**A**) Lung *Mtb*-infected monocyte and macrophage expression of *Spp1, Trem2,* and *Cxcl9*, (**B**) the resulting cell annotation, and (**C**) lung *Mtb*-infected monocyte and macrophage populations visualized separately for B6, *Nos2*^-/-^, *Acod1*^-/-^, and *Sp140*^-/-^ mice (n = 3 per genotype). (**D**) Quantification of the frequency of *Mtb*-infected IMs, *Spp1*^+^ IMs, and *Trem2*^+^ IMs among all IMs for B6, *Nos2*^-/-^, *Acod1*^-/-^, and *Sp140*^-/-^ mice (n = 3 per genotype). (**E**) Volcano plot of the differentially expressed genes in *Mtb*-infected *Spp1*^+^ IMs relative to *Trem2*^+^ IMs (combined analysis of cells from B6, *Nos2*^-/-^, *Acod1*^-/-^, and *Sp140*^-/-^ mice). (**F**) *Il1rn* expression measured by scRNA-seq of *Mtb*-infected myeloid cell populations in the lungs of B6, *Nos2*^-/-^, *Acod1*^-/-^, and *Sp140*^-/-^ mice (n = 3 per genotype). (**G**) Myeloid cell clustering and (**H**) volcano plot of the differentially expressed genes in *SPP1^+^* IMs relative to *TREM2^+^* IMs for cells from IgG control antibody-treated *Mtb*-infected non-human primate granulomas (data from 3 non-human primates, n = 12 granulomas; analysis of https://fairdomhub.org/studies/1134). (**I**) Myeloid cell clustering and (**J**) volcano plot of differentially expressed genes in *SPP1^+^* IMs relative to *TREM2^+^*IMs from 18F-FDG high (n = 5) and low (n = 6) portions of lung resections from *Mtb*-infected humans (analysis of GSE192483). (**K**) *IL1RN* expression by innate immune cells in FDG high and low signal portions of *Mtb*-infected human lungs as measured by scRNA-seq (analysis of GSE192483). (**L**) ROC curve depicting the ability of *IL1RN* expression to stratify human active *Mtb* patients (n = 151) and healthy controls (n = 98) in the TANDEM cohort (analysis of GSE114192). (**M**) ROC curve depicting the ability of *IL1RN* expression to stratify human active *Mtb* patients (n = 16) and LTBI individuals (n = 31) in the Berry South Africa cohort (analysis of GSE107992). Lungs were analyzed for the experiments depicted in (A) – (F) 25 days after *Mtb*-Wasabi infection and 6 weeks after *Mtb* infection for those depicted in (G) and (H). The bars in (D) represent the median. Parenthetical numbers in (L) and (M) indicate the 95% confidence interval. Statistical significance in (E), (F), (H), and (J) was calculated with the Wilcoxon Rank-Sum test with Bonferroni correction and in (D) by two-way ANOVA with Tukey’s multiple comparison test. ****p < 0.0001.

Given the conserved nature of IL-1Ra expression, we tested whether *IL1RN* expression could discriminate between active *Mtb* cases and healthy controls in RNA-seq of PBMCs from the TANDEM cohort^49^. We observed high sensitivity and specificity for *IL1RN* alone (AUC=0.84) (**Fig. 2L**), which was only slightly improved by including *IL1RN* in a gene signature containing additional known immunosuppressive molecules, such as *IL18BP*^58^, *CD274*^59^, and *IL10*^60^ (**Extended Data Fig. 5D**). *IL1RN* alone was also discriminatory when discerning active TB versus LTBI cases when applied to the Berry South Africa dataset^50^ (AUC=0.710; **Fig. 2M**). The high performance of *IL1RN* is consistent with prior work, as multiple published human TB signatures include *IL1RN*^8,41–43^. Analysis of *IL1RN* expression in the progressor population of the Adolescent Cohort Study^61^ identified that increased *IL1RN* expression was detectable ∼200 days before clinical TB diagnosis, this detection of elevated *IL1RN* expression during subclinical disease is in line with our hypothesis that IL-1Ra is a relatively early mechanism of disease exacerbation (**Extended Data Fig. 5E**). It is important to note that IFN response was the earliest detected transcriptional change in the Adolescent Cohort Study (observed ∼12-18 months before diagnosis; **Extended Data Fig. 5E**)^62^, suggesting that other factors initiate loss of bacterial control which may then be amplified by immunosuppressive molecules including IL-1Ra. As IL-1 signaling is regulated by both IL-1Ra and the decoy IL-1R2 receptor, we also examined *Il1r2* expression in our scRNA-seq datasets and found lower *Il1r2* expression in *Mtb*-infected animals relative to naïve controls, suggesting *Il1r2* does not contribute to *Mtb* susceptibility (**Extended Data Fig. 5A**). Taken together, our results indicate that myeloid cell expression of IL-1Ra is an important hallmark of TB disease that is conserved across mice, non-human primates, and humans.

### Markers distinguishing Spp1^+^ and Trem2^+^ interstitial macrophages

Given the strong correlation between *Spp1*^+^ macrophage differentiation, IL-1Ra expression, and *Mtb* disease progression, we sought to compare the *Spp1*^+^ IM response to *Mtb* infection among the *Mtb*-susceptible *Sp140*^-/-^, *Nos2*^-/-^, and *Acod1*^-/-^ mice. Our myeloid cell scRNA-seq dataset was used to determine potential markers for flow cytometric identification of *Spp1*^+^ IMs and *Trem2*^+^ IMs in infected mice. We found that *Itgax* (encoding CD11c) was upregulated on mature IMs, *Cd63* was preferentially expressed on *Trem2*^+^ IMs, and *Cd9* was preferentially expressed by *Spp1*^+^ IMs and was part of the core mouse *Spp1*^+^ signature (**Fig. 3A; Extended Data Fig. 4D, 6A, 6B**), enabling the analysis of these macrophage subsets by flow cytometry. CD166, also known as Alcam, was recently identified as a marker of human *SPP1*^+^ lung macrophages^63^, while *Trem2* and *C1q* are preferentially expressed by *TREM2*^+^ lung macrophages in mice, non-human primates, and humans by scRNA-seq (**Fig. 2E, 2H, 2J**). Using a gating strategy to distinguish IMs from AMs, we then gated on the mature PD-L1^hi^ CD11c^hi^ IMs and used CD9 and CD63 as markers for *Spp1*^+^ and *Trem2*^+^ IMs, respectively (**Fig. 3B, Extended Data Fig. 6C, 6D**). The PD-L1^hi^ CD11c^hi^ IMs were classified as mature macrophages due to their higher expression of CD64, PD-L1, and MHC II and lower expression of Ly6C relative to the PD-L1^low^ CD11c^low^ IMs (**Extended Data Fig. 6E, 6F**). CD9 and CD63 were validated as markers for these macrophage populations by assessing their staining for Alcam, Trem2, and C1q. As predicted, the CD9^hi^ *Spp1*^+^ IMs stained brighter for Alcam while the CD63^hi^ *Trem2*^+^ IMs stained brighter for Trem2 and C1q (**Fig. 3C**). We next assessed how *Mtb*-susceptibility impacts macrophage differentiation by comparing differentiation in B6, *Sp140*^-/-^, *Nos2*^-/-^, and *Acod1*^-/-^ mice.

**Figure 3.**
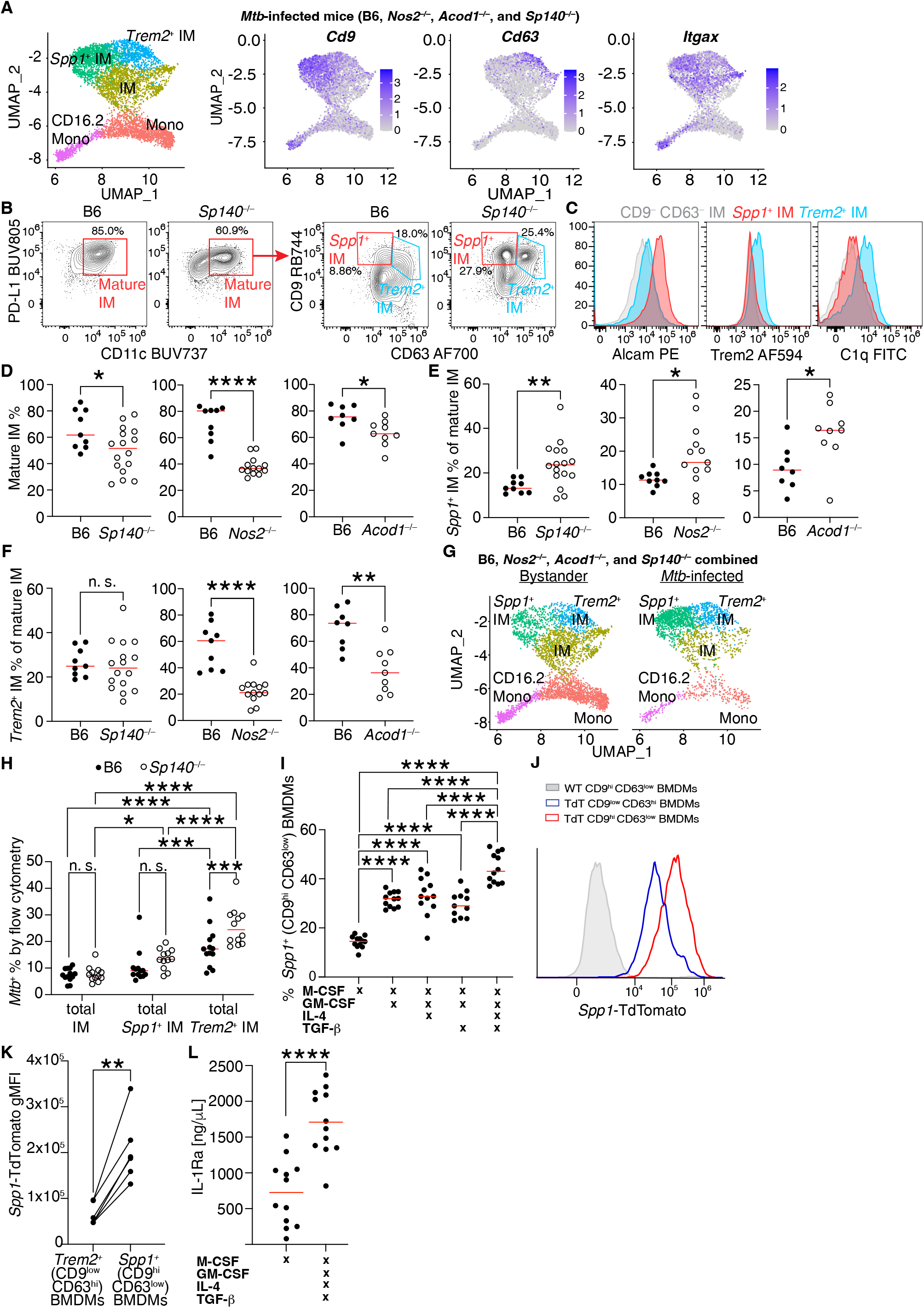
*Spp1*^+^ and *Trem2*^+^ macrophage differentiation is identifiable by flow cytometry, and *in vitro Spp1*^+^ macrophage differentiation correlates with IL-1 receptor antagonist production. (**A**) Lung *Mtb*-infected monocyte and macrophage expression of *Cd9, Cd63,* and *Itgax* plotted alongside annotated myeloid cell clusters for the *Mtb*-infected lung scRNA-seq dataset (combined analysis of cells from B6, *Nos2*^-/-^, *Acod1*^-/-^, and *Sp140*^-/-^ mice, n = 3 per genotype). (**B**) Gating scheme used to identify mature IMs (PD-L1^hi^ CD11c^hi^) as well as mature *Spp1*^+^ IMs (PD-L1^hi^ CD11c^hi^ CD9^hi^ CD63^low^) and mature *Trem2*^+^ IMs (PD-L1^hi^ CD11c^hi^ CD9^low^ CD63^hi^) in B6 and *Sp140*^-/-^ mice. (**C**) Representative histogram depicting *Spp1*^+^ IM (red), *Trem2*^+^ IM (blue), and IM (grey) expression of Alcam, Trem2, and C1q in the lungs of *Mtb*-infected mice. (**D**) Quantification of the frequency of the total mature IMs, (**E**) mature *Spp1*^+^ IMs, and (**F**) mature *Trem2*^+^ IMs in the lungs of *Mtb*-infected B6 (n = 9), *Sp140*^-/-^ (n = 16), *Nos2*^-/-^ (n = 13), and *Acod1*^-/-^ (n = 9) mice. (**G**) UMAP visualization comparing the frequency of the monocyte and macrophage clusters for bystander cells versus *Mtb*-infected cells (combined analysis of cells from B6, *Nos2*^-/-^, *Acod1*^-/-^, and *Sp140*^-/-^ mice, n = 3 per genotype). (**H**) Flow cytometric analysis of *Mtb* infection frequency among total (immature and mature) IM, *Spp1*^+^ IM, and *Trem*2^+^ IM in the lungs of B6 (n = 13) and *Sp140*^-/-^ mice (n = 12). (**I**) Frequency of BMDMs expressing the *Spp1*^+^ macrophage markers CD9^hi^ CD63^low^ after *in vitro* culture with M-CSF, GM-CSF, IL-4, and / or TGF-β (n=12). (**J**) Representative histogram and (**K**) quantification of the geometric mean fluorescence intensity (gMFI) of *Spp1*-TdTomato expression by *Spp1*^+^ (CD9^hi^ CD63^low^) and *Trem2*^+^ (CD9^low^ CD63^hi^) BMDMs from *Spp1-IRES-TdTomato* (TdT) and wild-type (WT) mice treated with M-CSF, GM-CSF, IL-4, and TGF-β (paired points indicate cells cultured in the same well; n=6). (**L**) IL1-Ra production by BMDMs cultured with M-CSF (n=12) or *Spp1*^+^ polarizing M-CSF, GM-CSF, IL-4, and TGF-β (n=12). Each dot represents an individual biological replicate, and colors indicate the differentiation condition. For (A) – (H), lungs were analyzed 24-28 days after *Mtb* or *Mtb*-Wasabi infection. The bars in (D) – (F), (H), (I), and (L) represent the median. Pooled data from two independent experiments are shown in (D) – (F), (H), (I), (K) and (L). Statistical significance in (H) and (I) was calculated by one-way ANOVA with Tukey’s multiple comparison test, in (D) – (F) by two-tailed t test, in (K) by two-tailed paired t test and in (L) by Pearson’s correlation coefficient (r). *p < 0.05, **p < 0.01, ***p < 0.001, ****p < 0.0001.

**Figure 4.**
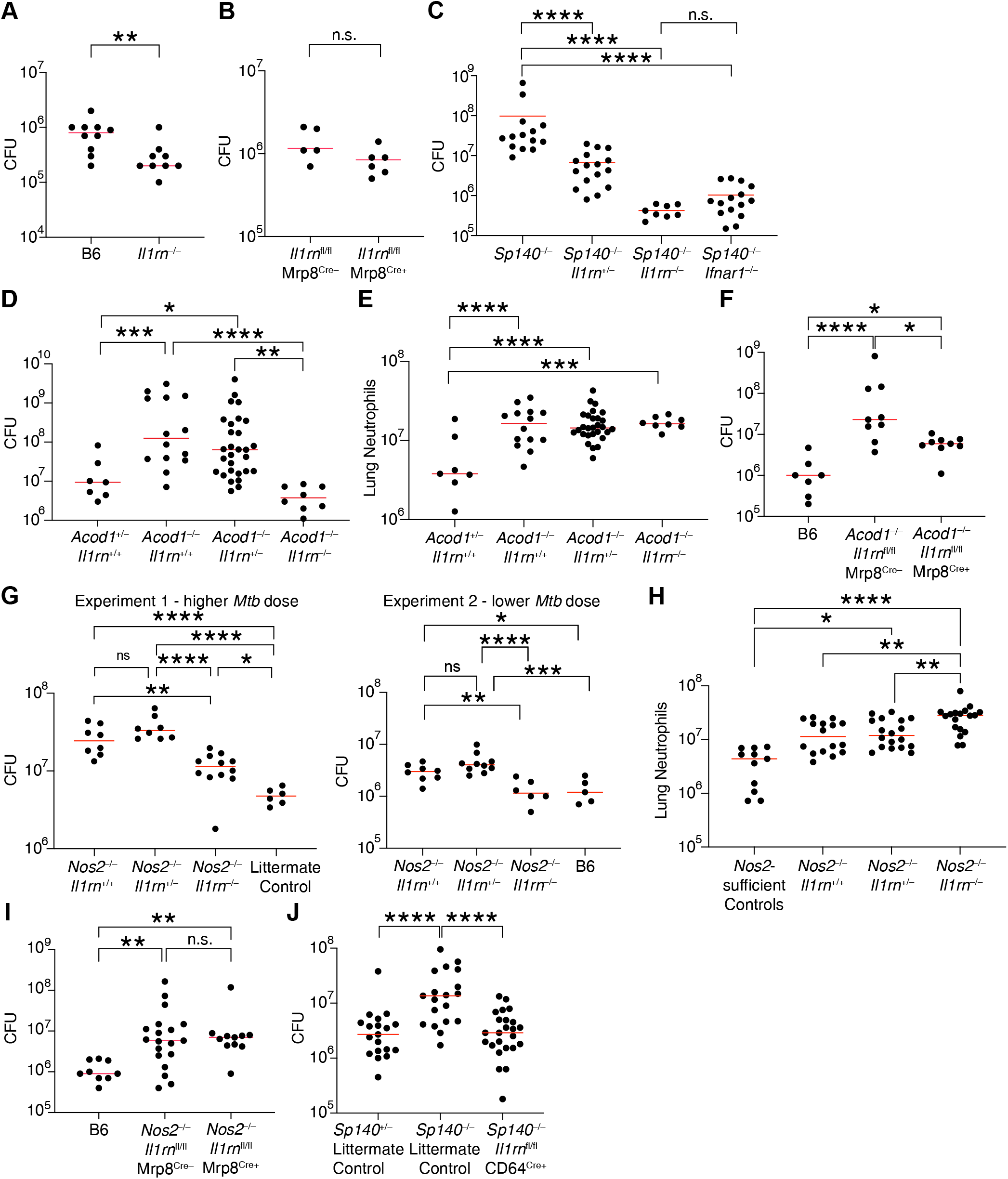
**Macrophage-derived IL-1Ra is a conserved driver of *Mtb* susceptibility.**(**A**) Lung bacterial burden in B6 (n = 10) and *Il1rn*^-/-^ mice (n = 9) following *Mtb* infection. (**B**) Lung bacterial burden in Mrp8^Cre-^ *Il1rn*^fl/fl^ (n = 5) and Mrp8^Cre+^ *Il1rn*^fl/fl^ mice (n = 6) following *Mtb* infection. (**C**) Lung bacterial burden in *Sp140*^-/-^ (n = 14), *Sp140*^-/-^ *Il1rn* ^+/-^ (n = 16), *Sp140*^-/-^ *Il1rn*^-/-^ (n = 8), and *Sp140*^-/-^ *Ifnar1*^-/-^ mice (n = 15) following *Mtb* infection. (**D**) Lung bacterial burden and (**E**) neutrophil numbers in *Acod1*^+/-^ *Il1rn*^+/+^ (n = 7), *Acod1*^-/-^ *Il1rn*^+/+^ (n = 14), *Acod1*^-/-^ *Il1rn*^+/-^ (n = 28), and *Acod1*^-/-^ *Il1rn*^-/-^ (n = 8) mice following *Mtb* infection. (**F**) Lung bacterial burden in B6 (n = 7), *Acod1*^-/-^ Mrp8^Cre-^ *Il1rn*^fl/fl^ (n = 9) and *Acod1*^-/-^ Mrp8^Cre+^ *Il1rn*^fl/fl^ mice (n = 9) following *Mtb* infection. (**G**) Lung bacterial burden and (**H**) neutrophil numbers in *Nos2*-sufficient controls (n = 11), *Nos2*^-/-^ *Il1rn*^+/+^ (n = 8-16), *Nos2*^-/-^ *Il1rn*^+/-^ (n = 8-18), *Nos2*^-/-^ *Il1rn*^-/-^ (n = 6-19), littermate control (n = 6) and B6 (n = 5) mice following *Mtb* infection. (**I**) Lung bacterial burden in B6 (n = 9), *Nos2*^-/-^ Mrp8^Cre-^ *Il1rn*^fl/fl^ (n = 19) and *Nos2*^-/-^ Mrp8^Cre+^ *Il1rn*^fl/fl^ mice (n = 11) following *Mtb* infection. (**J**) Lung *Mtb* burden in *Sp140*^+/-^ littermate control (n = 19), *Sp140*^-/-^ littermate control (including *Sp140*^-/-^ *Il1rn*^fl/fl^ *CD64^cre^*^-^, n = 19), and *Sp140*^-/-^ *Il1rn*^fl/fl^ *CD64^cre^*^+^ (n = 24) mice. Lungs were analyzed 24-28 days after *Mtb* or *Mtb*-Wasabi infection. The bars in (A) – (J) represent the median. Pooled data from two independent experiments are shown in (A) – (I) and three independent experiments in (J). Statistical significance in (A) and (B) was calculated with a two-tailed t test while in (C) - (J) it was calculated by one-way ANOVA with Tukey’s multiple comparison test. *p < 0.05, **p < 0.01, ***p < 0.001, ****p < 0.0001.

**Figure 5.**
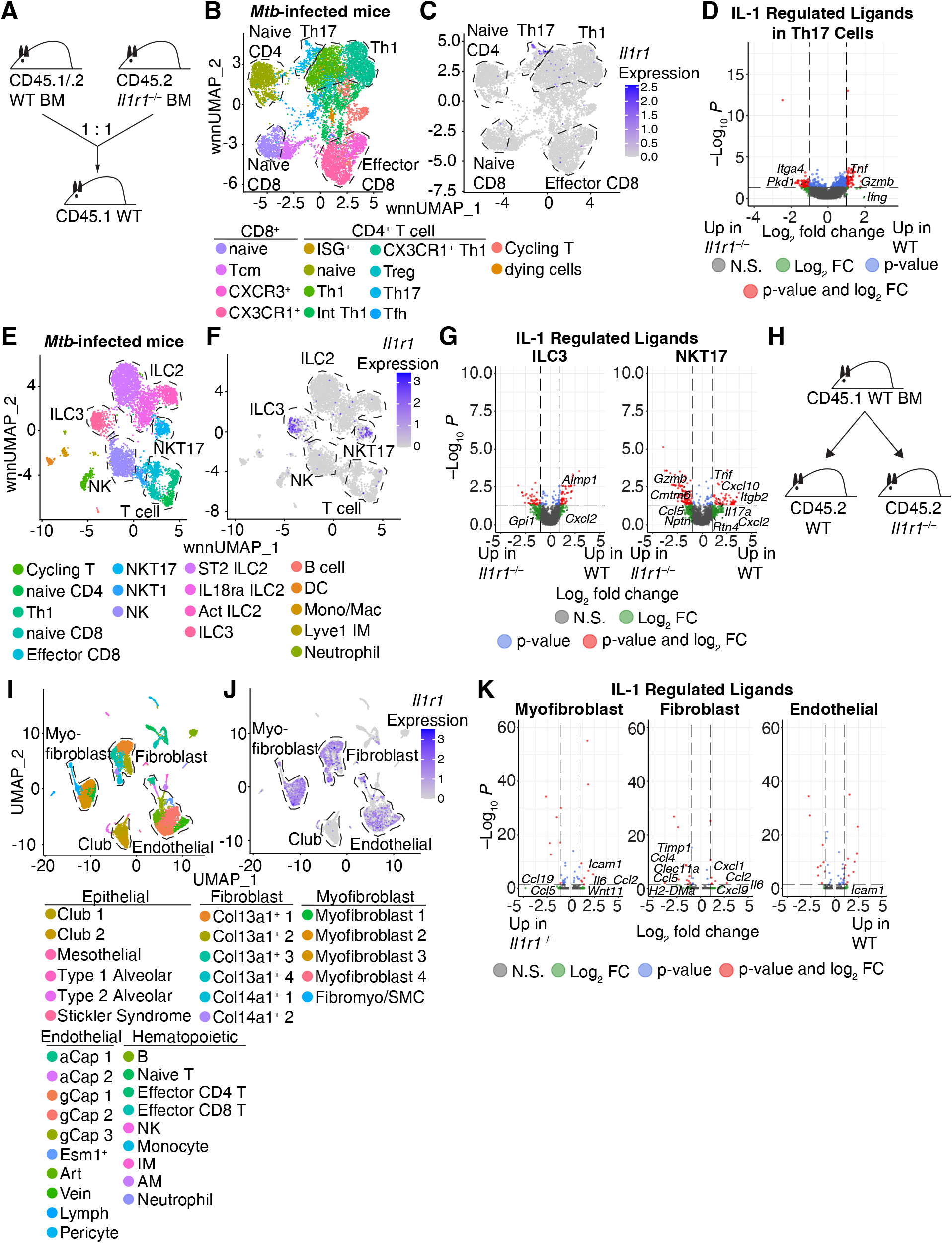
IL-1 enhances cytokine and chemokine expression by Th17, NKT17, and ILC3 as well as stromal cells. (**A**) Schematic of the BM chimera mice used in panels (B) – (G). (**B**) T cell populations and (**C**) their expression of *Il1r1* in the *Mtb*-infected lungs of the BM chimera mice (n = 3). (**D**) Volcano plot of the differentially expressed genes in the WT Th17 cells relative to *Il1r1*^-/-^ Th17 cells during *Mtb* infection. (**E**) Innate lymphoid cell populations and (**F**) their expression of *Il1r1* in the *Mtb*-infected lungs of the BM chimera mice (n = 3 with each sample consisting of 2-3 lungs pooled together). (**G**) Differentially expressed genes in the WT ILC3 and NKT17 cells relative to *Il1r1*^-/-^ cells during *Mtb* infection. (**H**) Schematic of the BM chimera mice used in panels (I) – (K). (**I**) Non-hematopoietic cell populations and (**J**) their expression of *Il1r1* in the *Mtb*-infected lungs of the BM chimera mice (n = 5 per genotype and infection status). (**K**) Differentially expressed genes in the WT myofibroblast, fibroblast, and endothelial cells relative to *Il1r1*^-/-^ cells during *Mtb* infection. Statistical significance in (D), (G), and (K) was calculated with the Wilcoxon Rank-Sum test with Bonferroni correction. *p < 0.05.

**Figure 6.**
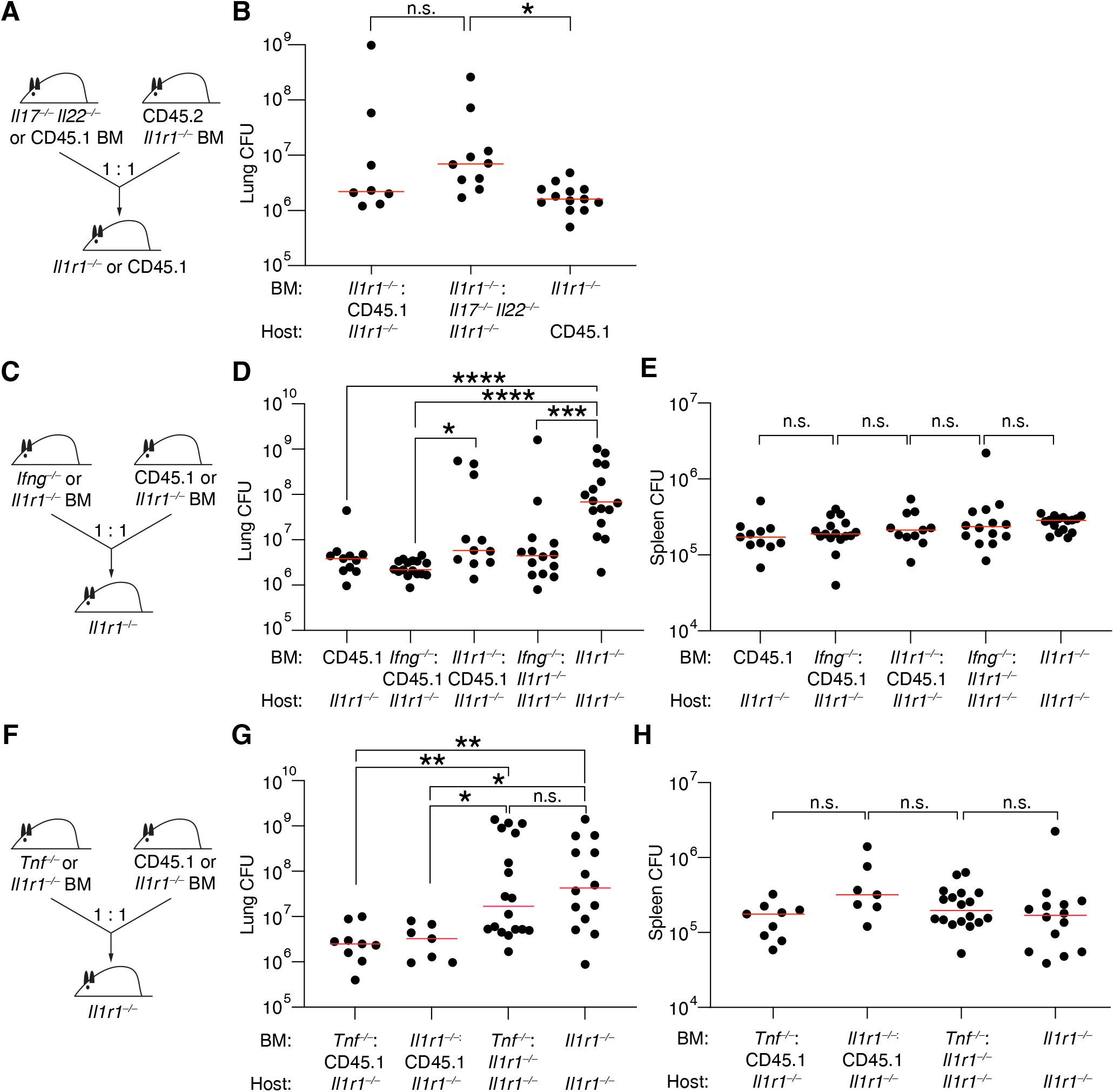
**IL-1 enhances *Mtb* control through amplification of TNF**. (**A**) Schematic of the BM chimera mice used in panel (B). (**B**) Lung *Mtb* burden in BM chimera mice that received *Il1r1*^-/-^: CD45.1 mixed BM (n = 8), *Il1r1*^-/-^: *Il17*^-/-^ *Il22*^-/-^ mixed BM (n = 10), or *Il1r1*^-/-^ BM (n = 13). (**C**) Schematic of the BM chimera mice used in panels (D) and (E). (**D**) Lung and (**E**) spleen *Mtb* burden in mixed BM chimera mice that received CD45.1 BM (n= 11), *Ifng*^-/-^: CD45.1 mixed BM (n = 16), *Il1r1*^-/-^: CD45.1 mixed BM (n = 11), *Ifng*^-/-^: *Il1r1*^-/-^ mixed BM (n = 14), and *Il1r1*^-/-^ BM (n = 16). **(F)** Schematic of the BM chimera mice used in panels (G) and (H)**. (G)** Lung and (**H**) spleen *Mtb* burden in mixed BM chimera mice that received *Tnf*^-/-^: CD45.1 mixed BM (n = 7), *Il1r1*^-/-^: CD45.1 mixed BM (n = 9), *Tnf*^-/-^: *Il1r1*^-/-^ mixed BM (n = 18), and *Il1r1*^-/-^: *Il1r1*^-/-^ BM (n = 14). Lungs were analyzed 25-26 days after *Mtb* or *Mtb*-Wasabi infection. The bars in (B), (D), (E), (F), and (G) represent the median. Pooled data from two independent experiments are shown in (B), (D), (E), (G), and (H). Statistical significance in (B), (D), (E), (G), and (H) was calculated by one-way ANOVA with Tukey’s multiple comparison test. *p < 0.05, **p < 0.01, ***p < 0.001, ****p < 0.0001.

Interestingly, all three susceptible mouse models displayed significant reductions in the frequency of mature IMs relative to B6 control mice, in line with the elevated monocyte recruitment observed in these animals following *Mtb* infection (**Fig. 3D, Extended Data Fig. 1A**). Of the mature IMs, there was a significant increase in *Spp1*^+^ macrophage frequency in all three susceptible models, while *Trem2*^+^ macrophage differentiation was reduced in *Nos2*^-/-^ and *Acod1*^-/-^ mice (**Fig. 3E, 3F**). We also quantified the total number of mature IMs, mature *Spp1*^+^ IMs, and mature *Trem2*^+^ IMs in these mice, and observed increases in mature IMs in the *Nos2*^-/-^ and *Acod1*^-/-^ mice but not *Sp140*^-/-^ mice relative to the B6 controls (**Extended Data Fig. 7A**). In line with the frequency data, we observed overall greater increases in mature *Spp1*^+^ IM numbers than in mature *Trem2*^+^ IM numbers, with the number of *Spp1*^+^ macrophages increased in *Nos2*^-/-^ and *Acod1*^-/-^ mice and, although not significant, ∼2-fold higher in *Sp140*^-/-^ mice relative to B6 animals (**Extended Data Fig. 7B, 7C**). These findings validate the scRNA-seq data and confirm that a shift from *Trem2*^+^ to *Spp1*^+^ IM differentiation correlates with loss of bacterial control (**Fig. 2D**).

As all three of the *Mtb*-susceptible mouse strains have increased bacterial burdens (**Fig. 1E, 1F, 1G**), we next asked whether *Mtb* infection directly drove *Spp1*^+^ IM differentiation. First, we reanalyzed (including re-annotation) a published scRNA-seq dataset of lung parenchymal monocytes and monocyte-derived macrophages from *Mtb*-infected C3H mice^64^, which are *Mtb*-susceptible and lack functional *Sp140*^14^. *Spp1*^+^ IMs were undetectable in naïve lungs and at day 10 of the infection, but a major population of the monocyte-derived macrophages by day 17 post-infection, once *Mtb* disseminated from AMs to monocyte-derived macrophages and neutrophils^65^ (**Extended Data Fig. 7D**). Given this association between *Mtb* spread to monocyte-derived macrophages and *Spp1*^+^ IM differentiation, we asked whether there was an association between direct *Mtb* infection and *Spp1*^+^ IM differentiation. While harboring *Mtb* correlated with an increase in *Spp1*^+^ and *Trem2*^+^ IMs by scRNA-seq (**Fig. 3G**), only the *Trem2*^+^ IMs exhibited a statistically significantly higher rate of infection by flow cytometry when comparing *Sp140*^-/-^ to B6 mice (**Fig. 3H**). On a per cell basis, *Mtb*-infected *Trem2*^+^ IM harbored more bacteria than *Spp1*^+^ IM in B6 mice, based on geometric mean fluorescence intensity (gMFI) of Wasabi expression in *Mtb*-Wasabi^+^ macrophages (**Extended Data Fig. 7E**). The *Trem2*^+^ IMs were also infected at a greater frequency than the *Spp1*^+^ IMs in B6 and *Sp140*^-/-^ mice (**Fig. 3H**). Additionally, most cells of either IM subset were uninfected, suggesting direct infection is not a major driver of *Spp1*^+^ IM differentiation. These data suggest that the inflammatory response to *Mtb* infection, rather than *Mtb* itself, promotes *Spp1*^+^ macrophage differentiation.

### *In vitro* differentiated *Spp1*^+^ macrophages secrete IL-1Ra

To test how components of the inflammatory response to *Mtb* infection promote *Spp1*^+^ macrophage differentiation, we turned to an *in vitro* macrophage differentiation approach. Lung GM-CSF production increases throughout the course of mouse *Mtb* infection^66,67^, and human monocytes have been shown to differentiate into SPP1 secreting CD9^+^ macrophages following *in vitro* culture with GM-CSF^68^. Therefore, we differentiated murine bone marrow cells isolated from wild-type (WT) or *Spp1*-TdTomato (TdT) reporter mice with M-CSF for 4 days, followed by stimulation with GM-CSF in combination with either IL-4, TGF-β, or both for 3 days to differentiate the cells into Spp1 expressing bone marrow-derived macrophages (BMDMs).

Macrophage differentiation was assessed by flow cytometry by classifying mature macrophages (MerTK^+^ CD64^+^ PD-L1^hi^ CD11c^hi^) as CD9^hi^ CD63^low^ *Spp1*^+^ macrophages and CD9^low^ CD63^hi^ *Trem2*^+^ macrophages (as gated in **Fig. 3B**). Interestingly, GM-CSF stimulation alone was sufficient to induce twice the proportion of CD9^hi^ CD63^low^ macrophages than M-CSF, with addition of IL-4 and TGF-β further enhancing *Spp1*^+^ (CD9^hi^ CD63^low^) macrophage differentiation (**Fig. 3I**). In support of CD9^hi^ CD63^low^ as markers of *Spp1*^+^ macrophages, CD9^hi^ CD63^low^ macrophages expressed more *Spp1* than CD9^low^ CD63^hi^ macrophages, as measured by *Spp1*-TdTomato gMFI (**Fig. 3J and 3K)**. Having validated the CD9^hi^ CD63^low^ gating scheme, we tested whether *Spp1*^+^ macrophage differentiation correlated with IL-1Ra secretion. In line with the high levels of *Il1rn* transcript expressed by *Spp1*^+^ macrophages in our scRNA-seq data, we observed that *Spp1*^+^ macrophage polarizing conditions (GM-CSF, IL-4, and TGF-β) significantly increased IL-1Ra secretion into the culture media relative to standard macrophage differentiation conditions (M-CSF; **Fig. 3L**). These findings are consistent with our *in vivo* experiments, suggesting that *Spp1*^+^ macrophages are the dominant source of IL-1Ra among monocytes and macrophages.

IL-1Ra expression has been associated with an M2-like phenotype in macrophages^69,70^, therefore we tested whether *Trem2*^+^ vs *Spp1*^+^ differentiation mapped onto the M1 vs M2 paradigm. Differentiation into an M1 polarized state was marked by iNOS expression, while an M2 polarized state was indicated by Arginase-1 (encoded by *Arg1*) expression. While Arg1 expression was largely restricted to the susceptible mouse models, iNOS and Arg1 were expressed by mature IMs, *Trem2*^+^ IMs, and *Spp1*^+^ IMs by scRNA-seq and flow cytometry demonstrating that *Trem2*^+^ vs *Spp1*^+^ differentiation is independent of canonical M1 vs M2 differentiation (**Extended Data Fig. 7F, 7G**).

### IL-1Ra is a conserved mediator of Mtb susceptibility

Next, we examined whether myeloid cell IL-1Ra expression plays a conserved role in amplifying *Mtb* disease. IL-1 is highly expressed in susceptible mouse strains^33^, and has been hypothesized to contribute to the susceptibility of *Nos2*^-/-^ mice to *Mtb*^13,33^. However, as we detected high levels of *Il1rn* expression in myeloid cells in *Sp140*^-/-^, *Nos2*^-/-^, and *Acod1*^-/-^ mice, we hypothesized that perhaps IL-1 signaling was impaired in these mice despite the high levels of IL-1 cytokine. We hypothesized that perhaps the susceptibility of the mice was due to a functional deficiency in IL-1 signaling. In support of the inhibitory role of *Il1rn* during *Mtb* infection, global deletion of *Il1rn* in B6 mice slightly reduced lung bacterial burden (**Fig. 4A**).

Neutrophils are major producers of *Il1rn* (**Fig. 2F**); however, neutrophils were not an obligatory source of *Il1rn*, as neutrophil-specific deletion of *Il1rn* using Mrp8^Cre^ *Il1rn*^fl/fl^ mice had no effect on bacterial burden on the B6 genetic background (**Fig. 4B**).

We previously found that *Mtb*-infected *Sst1*^S^ mice exhibit high levels of IL-1 and yet bacterial burdens in these mice are rescued by enhancement of IL-1 signaling via deletion of *Il1rn*^17^. Likewise, we found that even heterozygous deficiency in *Il1rn* promotes bacterial control in *Sp140*^-/-^ mice (**Fig. 4C**). However, as noted above (**Fig. 1**), *Sp140*^-/-^ and *Sst1*^S^ mice exhibit a strong type I IFN signature that is largely absent from *Nos2*^-/-^ and *Acod1*^-/-^ mice. Given that *Il1rn* is an interferon-stimulated gene, it was therefore of interest to test whether *Il1rn*-deficiency could also rescue bacterial burdens in *Nos2*^-/-^ and *Acod1*^-/-^ mice. We found that while deletion of one allele of *Il1rn* had no effect on *Mtb* control in *Acod1*^-/-^ mice, complete deletion of *Il1rn* was sufficient to restore wild-type levels of bacterial control at day 25 post-infection (**Fig. 4D**).

Interestingly, while bacterial burdens were reduced in *Acod1*^-/-^ *Il1rn*^-/-^ mice to levels seen in WT B6 mice, lung neutrophil numbers were still elevated in *Acod1*^-/-^ *Il1rn*^-/-^ mice (**Fig. 4E**). This result suggests that production of IL-1Ra may be an important mechanism by which neutrophils drive *Mtb* susceptibility. However, neutrophil-specific deletion of *Il1rn* in *Acod1*^-/-^ mice only slightly improved bacterial restriction, suggesting that while neutrophils may be a source of *Il1rn* during *Mtb* infection, they may not be the sole or dominant source (**Fig. 4F**). In two experiments with a higher and lower dose of *Mtb* inoculum, we also observed partial or complete rescue of bacterial burdens in *Nos2*^-/-^ *Il1rn*^-/-^ mice as compared to *Nos2*^-/-^ mice (**Fig. 4G**). Similar to the *Acod1*^-/-^ mice, no effect of *Ilr1rn* heterozygosity was observed, and the improved bacterial control in *Nos2*^-/-^ *Il1rn*^-/-^ mice did not coincide with a decrease in the number of lung neutrophils (**Fig. 4H**). Despite the increase in neutrophils, neutrophil expression of *Il1rn* did not seem to be a major driver of the *Mtb* susceptibility for *Nos2*^-/-^ mice, as neutrophil-specific deletion of *Il1rn* had no effect on bacterial control in these mice (**Fig. 4I**). These results demonstrate a conserved role for IL-1Ra in elevating *Mtb* burdens in three different mouse models at relatively early infection timepoints, but do not rule out additional roles for IL-1Ra in suppressing immunopathology at later timepoints during TB.

While neutrophil expression of IL-1Ra is constitutive (including in resistant B6 mice), it is induced in IMs (**Extended Data Fig. 5A, 5B**). Indeed, *Spp1*^+^ IMs exhibited the highest expression levels of *Il1rn* among monocytes and macrophages and *Spp1*^+^ IMs in all three susceptible mouse models expressed elevated levels of *Il1rn* relative to wild-type B6 controls (**Fig. 2F**). Additionally, macrophages localize closer to *Mtb* than neutrophils in all three susceptible mouse models (**Extended Data Fig. 1G**). We therefore hypothesized that macrophage production of IL-1Ra could account for the conserved role of IL-1Ra in inhibiting bacterial control in *Mtb*-susceptible mice. To delete *Il1rn* in macrophages, we used CD64-Cre mice, which express Cre specifically in macrophages, at least in naïve animals^71^. Deletion of *Il1rn* in CD64^Cre^ *Il1rn*^fl/fl^ mice^71,72^ was able to rescue *Mtb* susceptibility of *Sp140*^-/-^ mice (**Fig. 4J**). Together, these data reveal a conserved, myeloid cell role for IL-1Ra expression in exacerbating *Mtb* disease.

### IL-1-enhanced cytokine and chemokine production promotes Mtb control

Given the conserved role for IL-1Ra in exacerbating *Mtb* infection, we aimed to better define how IL-1 mediates protection against *Mtb*. We first examined IL-1R expression on myeloid cells, as they are the primary reservoir of *Mtb*, and found little to no expression of IL-1R in myeloid cells from naïve and *Mtb*-infected B6, *Sp140*^-/-^, *Nos2*^-/-^, and *Acod1*^-/-^ mice (**Extended Data Fig. 8A, 8B**). Consistent with a previous study^23^, we also demonstrated that IL-1R-deficient macrophages do not exhibit a cell-intrinsic defect in bacterial control in mixed bone marrow (BM) chimera mice (**Extended Data Fig. 8C, 8D, 8E**). It thus appears that IL-1 mediates its protective effect by signaling in uninfected bystander cells that then promote myeloid cell *Mtb* control. We therefore sought to identify the bystander cells that express IL-1R and the genes regulated by IL-1 in these cells to determine the mechanism by which IL-1 aids in bacterial restriction.

As IL-1 signaling in either hematopoietic or non-hematopoietic cells is sufficient for *Mtb* control during the initial few weeks of infection^23,39^, we sought to identify potential IL-1 responders in both compartments. Due to the inhibition of IL-1 signaling by high expression of IL-1Ra in the *Mtb*-susceptible mouse models, we used resistant B6 mice for examining IL-1 function. To ascertain the cell autonomous effects of IL-1, we generated mixed BM chimera mice with a 1:1 mix of WT and *Il1r1*^-/-^ cells that were infected with *Mtb* following immune reconstitution (**Fig. 5A**). Among hematopoietic cells, IL-1 promotes cytokine expression in CD4^+^ T cells^73^ and innate lymphoid cells (ILCs)^74,75^. In T cells, *Il1r1* was primarily expressed in T helper 17 cells (Th17) with some expression in Type 1 helper cells (Th1; **Fig. 5B, 5C, Extended Data Fig. 9A**). Differential scRNA-seq gene expression profiling comparing WT and *Il1r1*^-/-^ Th17 cells identified *Tnf*, *Gzmb*, and *Ifng* as genes with at least 2-fold greater expression in WT cells that were annotated as potential ligands in the NicheNet database^76^ (**Fig. 5D**). Type 3 ILC (ILC3) and IL-17 producing natural killer T (NKT17) cells were the major *Il1r1-*expressing innate lymphoid cell populations in *Mtb*-infected lungs, with differential gene expression analysis identifying *Aimp1* and *Cxcl2* as potential IL-1-induced ligands produced by ILC3s; and *Tnf*, *Cxcl10*, *Itgb2*, *Il17a*, *Cxcl2*, and *Rtn4* as potential IL-1-induced ligands produced by NKT17 cells (**Fig. 5E, 5F, 5G, Extended Data Fig. 9B**).

Regarding potential non-hematopoietic sensors of IL-1, we first examined sensory neurons as they highly express IL-1R^77^, and IL-1 signaling in neurons recruits myeloid cells^78^. As IL-1 signaling in hematopoietic cells is sufficient for initial resistance to *Mtb* infection^23,39^, we generated BM chimera mice that lacked hematopoietic IL-1R and either had IL-1 receptor in sensory neurons (CD45.1 and Adv^+/+^ *Il1r1*^fl/fl^ recipient mice) or not (Adv^cre/+^ *Il1r1*^fl/fl^ recipient mice). Sensory neuron-specific deletion of *Il1r1* had no effect on *Mtb* control, suggesting that other non-hematopoietic cells mediate the protective effect of IL-1 against *Mtb* (**Extended Data Fig. 8F**, **8G**). We thus performed scRNA-seq on BM chimera mice generated by transferring WT BM into lethally irradiated WT and *Il1r1*^-/-^ hosts and then infected the chimeric mice with *Mtb* following immune reconstitution (**Fig. 5H**). Lineage markers used to annotate a scRNA-seq dataset of stromal cells in normal and hyperoxia-damaged murine lungs^79^ were used to identify cell populations in our stromal cell scRNA-seq dataset (**Extended Data Fig. 9C**). Based on this analysis, *Il1r1* was widely expressed in stromal cells, including in myofibroblasts, fibroblasts, and endothelial cells (**Fig. 5I, 5J**). Differential expression analysis of IL-1-regulated potential ligands, as annotated in NicheNet’s database^76^, identified *Icam1*, *Il6*, *Ccl2*, and *Wnt11* in myofibroblasts, *Cxcl1*, *Ccl2*, *Il6*, and *Cxcl9* in fibroblasts, and *Icam1* in endothelial cells as IL-1-induced potential ligands (**Fig. 5K**).

The most likely candidates for the protective effect of IL-1 signaling were then identified using NicheNet^76^ (**Extended Data Fig. 10A**). For this analysis, we focused on IMs as the receiver cells for the IL-1-regulated ligands, as IMs are known to harbor *Mtb* and control the infection^80^. According to the NicheNet ligand-receptor database, the receptors for IFN-γ, TNF, IL-6, CCL2, and IL-17 were expressed by IMs (**Extended Data Fig. 10B**). While IFN-γ and TNF have well-established roles in *Mtb* control^3–5^, the contribution of IL-17 has been appreciated more recently^81–84^. IL-22 is another cytokine important for bacterial control that is often co-produced with IL-17^85^. Thus, we tested whether IL-1-induced enhancement of IL-17/22 production was important in IL-1-mediated *Mtb* restriction. This hypothesis was tested by generating mixed BM chimera mice containing BM from *Il17*^-/-^ *Il22*^-/-^ and *Il1r1*^-/-^ mice mixed at a 1:1 ratio and injected into lethally irradiated *Il1r1*^-/-^ recipient animals as well as control mice

(**Fig. 6A**). While the *Il17*^-/-^ *Il22*^-/-^: *Il1r1*^-/-^ BM chimera mice contain cells that can respond to IL-1, as well as cells that can produce IL-17/22, no cell will be able express IL-17/22 in response to IL-1. Thus, these chimeras specifically eliminate the ability of IL-1 to amplify IL-17/22 production. The *Il17*^-/-^ *Il22*^-/-^: *Il1r1*^-/-^ BM chimera mice were only slightly more susceptible than control animals, suggesting that an IL-1-mediated increase in IL-17/22 expression is not necessary for IL-1-driven protection (**Fig. 6B**).

Therefore, we sought to identify which other potential ligands were the most likely drivers of IL-1-regulated *Mtb* restriction by IMs. We identified genes upregulated in activated IMs, and NicheNet^76^ was then used to calculate the likelihood for each of the potential ligands to induce the transcriptional changes observed in the activated IMs. IFN-γ and TNF had the highest scores indicating they are the most likely candidates for how IL-1 protects against *Mtb*, while IL-17 had one of the lowest scores, in line with our experimental data (**Fig. 6B, Extended Data Fig. 10C**). Thus, we tested whether IFN-γ or TNF production is important for IL-1-mediated *Mtb* restriction by generating mixed BM chimera mice containing BM from *Ifng*^-/-^ or *Tnf*^-/-^ mice and *Il1r1*^-/-^ mice mixed at a 1:1 ratio and administered to lethally irradiated *Il1r1*^-/-^ recipients. While the *Ifng*^-/-^: *Il1r1*^-/-^ BM chimera mice contain cells that can respond to IL-1, as well as cells that can produce IFN-γ, no cell will be able to express IFN-γ in response to IL-1 (**Fig. 6C**). Thus, these chimeras specifically eliminate the ability of IL-1 to amplify IFN-γ production. Surprisingly, no significant increases in bacterial loads were observed in the lungs or spleen of *Ifng*^-/-^: *Il1r1*^-/-^ BM chimera mice relative to the control animals, indicating that IL-1-mediated enhancement of IFN-γ expression is dispensable for IL-1-driven protection (**Fig. 6D, 6E**). Interestingly, mice that only received *Il1r1*^-/-^ BM were susceptible to infection with elevated lung bacterial burden but had no increase in splenic burden, suggesting IL-1 does not play a major role in controlling *Mtb* dissemination in line with a previous report^86^.

Using the BM chimera mice that were reconstituted with BM from *Tnf*^-/-^ and *Il1r1*^-/-^ mice, we tested the role of IL-1 amplified TNF production on *Mtb* control (**Fig. 6F**). Similar to the previous BM chimera mice, these mice eliminate TNF production in response to IL-1. Following *Mtb* infection, we found these *Tnf*^-/-^: *Il1r1*^-/-^ BM chimera mice had increased pulmonary bacterial burden and were as susceptible as the *Il1r1*^-/-^ group (**Fig. 6G**). However, the splenic burden was not increased, which aligns with our previous observations (**Fig. 6H**). The *Tnf*^-/-^: *Il1r1*^-/-^ BM chimera mice also had increased total numbers of innate and adaptive immune cells, suggesting that the increased bacterial load is not due to impaired immune cell trafficking to the lungs (**Extended Data Fig. 10D, 10E**). Taken together, these results demonstrate that IL-1 mediates pulmonary control of bacterial burden by amplifying TNF production.

## Discussion

*Mtb* infection induces chronic inflammation through type I/II IFN induction up to 12-18 months prior to disease diagnosis^27,62^. While aspects of this inflammatory state—such as the IFN-γ response^3,4^—promote bacterial control, other components, such as the type I IFN response, exacerbate *Mtb* disease progression^8,15,22,87^. Our results suggest that the chronic inflammatory nature of *Mtb* infection may hinder bacterial control by promoting a state of immunosuppression that is conserved across susceptible mouse models, non-human primates and humans.

Previous work on immunosuppression during *Mtb* infection has identified PD-1, IDO1, TGFβ, and IL-10 as potential pathways that inhibit productive host responses^88–90^. However, the impact of immune suppression by myeloid cells on *Mtb* control is unclear, and can vary between models. For example, targeting the PD-1 pathway can exacerbate TB disease^91,92^, while targeting IDO1 can have no effect or promote *Mtb* control^93–95^, and deletion or blockade of IL-10 has varying effects in mice depending on the genetic background and infection conditions^11,96–100^.

Therefore, we applied a comparative approach to re-examine myeloid cell-mediated immunosuppression during *Mtb* infection. Gene signatures consisting of *Mtb*-induced genes expressed by myeloid cells in the lungs of *Mtb*-susceptible *Sp140*^-/-^, *Nos2*^-/-^, and *Acod1*^-/-^ mouse lungs^13–15,21^ significantly outperform wild-type B6 mouse signatures in discriminating humans with TB disease from healthy controls and LTBI individuals, suggesting susceptible mice better mimic aspects of human disease. Due to the high predictive strength of all three models’ gene signatures, it is likely that there are commonalities among the mouse models that are responsible for their discriminatory ability. We initially hypothesized that type I IFN was a likely candidate for a shared response to *Mtb* infection, as viral infections and the anti-viral type I IFN response correlate with the disease severity in humans^8,62,101–103^. Type I IFNs exacerbate *Mtb* infection in B6 mice^36,104–109^ and *Ifnar1* deletion rescues bacterial control in *Sst1*^S^ and *Sp140*-deficient mice^14,15,17^. To our surprise, however, a type I IFN signature was not conserved across susceptible mouse strains, and type I IFN receptor deletion had little to no effect on bacterial control in *Nos2*^-/-^ and *Acod1*^-/-^ mice, indicating that other pathways drive their *Mtb* susceptibility.

All three tested *Mtb*-susceptible mouse models exhibited an increase in *Spp1*^+^ macrophages. Interestingly, *Mtb*-restrictive B6 mice largely lacked *Spp1*^+^ macrophages, and thus, *Spp1*^+^ macrophage differentiation correlated with *Mtb* susceptibility. This result is also observed in multiple human diseases, suggesting that chronic inflammation is a driver of SPP1^+^ macrophage differentiation. For example, increased SPP1^+^ macrophage differentiation has recently been identified in the bronchoalveolar lavage fluid of active TB patients^110^, in individuals with idiopathic pulmonary fibrosis^63^, individuals with active Rheumatoid arthritis^111^, and in the lungs of individuals with severe COVID-19^111^. Additionally, SPP1 plasma levels correlate with human active TB disease^112^. Macrophage differentiation into the SPP1^+^ state also strongly correlates with tumor progression in humans^54^. Unlike this cancer study, we did not observe a negative correlation between macrophage expression of *Spp1* and *Cxcl9*. IFN-γ potently induces *Cxcl9* expression^15^, and we observed conserved, high levels of IFN-γ signaling in B6 and *Mtb*-susceptible mice. Therefore, it is possible that the inverse relationship between *Spp1* and *Cxcl9* expression in human head and neck cancer patients^54^ is masked by the strong IFN-γ response present during *Mtb* infection.

Despite the strong correlation between chronic inflammation and SPP1^+^ macrophage differentiation, the specific pathways promoting their differentiation and mediating their function remain underexplored. Therefore, we established CD9 and CD63 as flow cytometry markers to distinguish mouse *Spp1*^+^ from *Trem2*^+^ macrophages, which were validated with SPP1 reporter mice and an *in vitro* differentiation approach we developed. Using these markers, we demonstrated that direct *Mtb* infection is not required for *Spp1*^+^ differentiation *in vivo*. Instead, our *in vitro* system identified GM-CSF as a key cytokine for *Spp1*^+^ macrophage differentiation. These findings are consistent with a previous study showing that GM-CSF, together with TGF-β or IL-17, drives SPP1 and CD9 expression by human macrophages^68^. However, our *in vitro* system resulted in a mixed culture with only ∼45% of the macrophages adopting the *Spp1*^+^ fate, indicating that additional signals likely contribute to *Spp1*^+^ macrophage differentiation. Using the CD9 and CD63 gating scheme, we were also able to demonstrate that *Spp1*^+^ versus *Trem2*^+^ differentiation does not map onto the classic M1 versus M2 paradigm with both populations expressing canonical markers of M1 and M2 differentiation (e.g. iNOS and Arg1). These results are consistent with a growing appreciation that the M1/M2 paradigm, while valuable, is likely an oversimplification of the complexity of myeloid cell differentiation *in vivo*^113^.

Regarding *Spp1*^+^ macrophage function, our scRNA-seq analysis indicated that *Spp1*^+^ macrophages express *Il1rn*, and our *in vitro* differentiation approach confirmed a strong correlation between *Spp1*^+^ macrophage differentiation and IL-1Ra secretion. IL-1Ra inhibits IL-1 signaling. While IL-1-deficient mice are clearly susceptible to *Mtb*^23,36–40^ and anecdotal evidence in humans reports *Mtb* reactivation following IL-1Ra (Anakinra) treatment^114^, many susceptible mouse models exhibit elevated levels of IL-1^17,33^ leading some to propose that IL-1-driven neutrophilic inflammation actually exacerbates *Mtb*^13,29^. Here we propose that high levels of IL-1 cytokine do not necessarily imply high levels of IL-1 signaling. Indeed, we observed that increasing IL-1 signaling by global deletion of IL-1Ra improves bacterial control in *Sp140*^-/-^, *Nos2*^-/-^, and *Acod1*^-/-^ mice. The finding that global *Il1rn* deletion partially rescues *Nos2*^-/-^ mouse susceptibility was highly unexpected given prior claims that excessive IL-1 signaling is detrimental during *Mtb* infection^35^, particularly following the onset of the adaptive immune response in *Nos2*^-/-^ mice^13,33^. Thus, our results suggest that the initial lack of bacterial control in highly inflamed *Mtb*-susceptible mouse models is primarily due to insufficient rather than excessive IL-1 signaling. Importantly, our results do not exclude the possibility that excessive IL-1 signaling at later time points might have additional detrimental effects^115^. This balance is exhibited in our previous work in which homozygous deletion of *Il1rn* completely rescued bacterial control in *Sst1*^S^ mice 4 weeks post-infection but resulted in only a minor improvement in survival, while deletion of a single allele of *Il1rn* nearly phenocopied the survival of *Mtb*-restrictive B6 mice^17^.

In addition to macrophages as a source of IL-1Ra, our scRNA-seq data demonstrated that *Il1rn* was constitutively expressed by neutrophils. While neutrophil-specific deletion of *Il1rn* resulted in little to no improvement in bacterial control in *Acod1*^-/-^ and *Nos2*^-/-^ mice, we found that expression of IL-1Ra by CD64^+^ cells, such as macrophages, is specifically required for *Mtb* control in *Sp140*^-/-^ mice. These results recapitulate findings in *Candida* infection, during which neutrophils secreted considerably less IL-1Ra than monocytes upon stimulation and macrophage-specific deletion of *Il1rn* restored control of *Candida* infection^116^. Additionally, we identified that macrophages were located significantly closer to *Mtb* than neutrophils in the diseased portion of the lungs of all three susceptible mouse strains. This result correlates with human disease, as SPP1^+^ macrophages were found to reside near human TB granulomas^117^, and macrophages are one of the primary IL-1Ra-expressing cell types in human *Mtb* granulomas^118^. Together, these results suggest that low IL-1Ra secretion by neutrophils and differences in positioning within *Mtb* granulomas may contribute to the greater impact of macrophage-derived IL-1Ra on bacterial control.

Importantly, our study focused on assessing bacterial burdens at a relatively early time point after infection (4 weeks), as elevated *IL1RN* transcript was detectable ∼200 days prior to clinical diagnosis of TB in the Adolescent Cohort Study. However, *IL1RN* is not the initial trigger for loss of bacterial control as type I interferon responses are detectable 12-18 months prior to diagnosis in the same cohort^62^. A recent study of proteomic biomarker analysis in two human cohorts^119^ recapitulated this observation, with the type I interferon-driven CXCL10^15^ discriminating symptomatic and asymptomatic TB-infected individuals from healthy controls, while IL-1RA demonstrated strong performance in symptomatic individuals, but weaker, although still positive, performance in asymptomatic cases. Therefore, *Spp1*^+^ macrophage differentiation and their IL-1Ra production is likely not the first step in TB progression but instead part of a positive feedback loop that accelerates progression, potentially in only a subset of human patients.

The mechanism by which IL-1 exerts its protective effects remains the subject of intensive study. In line with the limited expression of *Il1r1* by infected myeloid cells in our scRNA-seq dataset, IL-1 has been proposed to act on uninfected bystander cells that then promote myeloid cell-mediated *Mtb* control^23,39^. IL-1 signaling in either hematopoietic or non-hematopoietic bystander cells is sufficient to mediate protection during the initial weeks of infection^23,39^. Yet the mechanism for IL-1-mediated protection is not fully established, with eicosanoid regulation and synergistic interactions between IL-1 and TNF being two potential mechanisms^36,39^. We identified multiple IL-1-induced cytokines and chemokines as potential activators of *Mtb*-harboring IMs, with the most likely being IFN-γ and TNF as they are known to play a key role in *Mtb* restriction and displayed the strongest signal in our macrophage NicheNet analysis^3–5^.

As synergy between IL-1 and TNF has been proposed as a mechanism for *Mtb* control^39^ but has not been directly assessed, we sought to test the role of IL-1 amplification of IFN-γ and TNF on *Mtb* control. Surprisingly, we observed that IL-1-mediated enhancement of IFN-γ is dispensable for protection against *Mtb*. This observation could be due to a difference in the sites of action for IFN-γ and IL-1. Indeed, Sakai et al.^120^ reported that IFN-γ plays a minimal role in controlling *Mtb* in the lungs, and instead is important in limiting dissemination to other organs, including the spleen. Conversely, our data indicate that IL-1 does not significantly regulate splenic bacterial burden. These tissue-specific differences in the primary action of these cytokines may explain the lack of an observed synergistic effect between IL-1 and IFN-γ in our experiments. In contrast, we observed that IL-1-enhanced TNF production is required for pulmonary *Mtb* control. As IL-1 signaling in CD4^+^ T cells directly promotes cytokine production^73^, it is likely that IL-1 directly enhances TNF production by T cells that then acts on lung *Mtb*-harboring IMs to promote their restriction of the infection. The site-specific differences in cytokine function we observed in our mouse studies resemble human *Mtb* infection. TNF-deficient individuals experienced recurrent pulmonary TB^121^, while individuals with mutations in IFN-γ signaling commonly present with disseminated Mycobacterial disease^122^, including disseminated TB^123,124^.

Regarding the role of IL-1 on non-hematopoietic cells, IL-1 acts on type II alveolar epithelial cells by inducing GM-CSF during *Legionella pneumophila* infection^125^, another cytokine that activates macrophages with an established contribution to *Mtb* control^67,126,127^. Our stromal scRNA-seq dataset had limited coverage of epithelial cells, so we were unable to observe whether IL-1 also induces GM-CSF expression in type II alveolar cells during *Mtb* infection. However, it is likely that this mechanism is conserved between *Legionella* and *Mtb* infections given the critical role played by GM-CSF and stromal cell IL-1 signaling in both infections.

In summary, our data support a model in which initial loss of bacterial control, whether due to a type I IFN response, *Nos2*-, or *Acod1*-deficiency, results in myeloid cell influx into *Mtb*-infected lungs. The incoming monocytes differentiate into IMs that adopt the *Spp1*^+^ activation state in animals with heightened inflammation, which correlates with the upregulation of IL-1Ra. This high level of IL-1Ra expression hinders critical IL-1 signaling in hematopoietic and non-hematopoietic cells, reducing the production of *Mtb*-restrictive cytokines, including TNF production by the hematopoietic cells. The IL-1Ra-mediated blockade of IL-1 thus generates a positive feedback loop promoting bacterial replication in which *Mtb*-harboring myeloid cells are unable to restrict the bacteria. Enhanced bacterial replication leads to the influx of yet more myeloid cells and further increases IL-1Ra levels. While intervention abolishing this feedback loop (e.g. IL-1Ra blockade) may temporarily promote bacterial control, the long-term cost of such an intervention would likely be an increased risk of immunopathology as this intervention would not address the initial insult that rendered the individual susceptible to TB, such as a strong type I IFN response. In this study, we provide molecular evidence for a feedforward loop of disease exacerbation by elucidating key molecular and cellular players that orchestrate the shift from *Mtb* restriction to susceptibility.

## Acknowledgments

We thank members of the Philips, Vance, Barton, Stanley, and Cox laboratories for advice and discussions; the Stanley and Cox labs, and in particular D. Fox, for the shared operation of the UC Berkeley BSL-3 and ABSL-3 labs; M. Cain for the operation of the WashU Med ABSL-3 labs; S. Gupta and P. Khatri for advice on analysis of human cohorts; B. Malissen, S. Khader, and Y. Belkaid for mice; W. Hasenpflug, P. Dietzen, R. Chavez, J. Morales, and J. Rodriguez for technical assistance; A. Valeros, H. Nolla, and K. Heydari of the UC Berkeley Cancer Research Laboratory Flow Cytometry facility for assistance with flow cytometry; the Alvin J. Siteman Cancer Center at WashU Medicine and Barnes-Jewish Hospital in St. Louis, MO., for the use of the Siteman Flow Cytometry core, which is supported in part by an NCI Cancer Center Support Grant #P30 CA091842; H. Aaron and F. Ives of the UC Berkeley Cancer Research Laboratory Molecular Imaging Center (RRID:SCR_017852) for assistance with confocal microscopy (supported by the Helen Wills Neuroscience Institute). Model figures were created with BioRender.com. D.I.K. is supported by NIH grants F32HL158250 and K22AI179939. C.G. is supported by the Swiss National Science Foundation (310030B_201269) and the Rheumasearch Foundation. S.S. is supported by the NIH grant R01AI175614. R.E.V. is an Investigator of the Howard Hughes Medical Institute and is supported by NIH grants AI075039, AI063302, and AI155634.

## Author Contributions

Conceptualization: D.I.K. and R.E.V. Investigation: D.I.K., O.V.L., E.P., S.H.B., and D.X.J. Data analysis: D.I.K., O.V.L., E.P., S.H.B., and B.A.R. Methodology: S.S. and M.M. Resources: C.G. and D.L.J. Writing: D.I.K., E.P., S.H.B., and R.E.V. with input from all co-authors. Funding acquisition: D.I.K. and R.E.V. Supervision: D.I.K. and R.E.V.

## Materials and Methods

### Mice

Following University of California, Berkeley and WashU Medicine Institutional Animal Care and Use Committee regulatory standards, mice were housed at 23°C with a 12-hour light-dark cycle and maintained under specific pathogen-free conditions. Male and female mice were used for experiments. Mice were age-matched with littermate controls when possible and 6-12 weeks old at the start of the infections. C57BL/6 (B6), B6.129S2-Ifnar1^tm1Agt^/Mmjax (*Ifnar1*^-/-^)^128^, B6.SJL-*Ptprc^a^ Pepc^b^*/BoyJ (CD45.1)^129^, B6.129S7-*Il1r1^tm1Imx^*/J (*Il1r1*^-/-^)^130^, B6.129(Cg)-*Il1r1^tm1.1Rbl^*/J (*Il1r1*^fl/fl^)^131^, B6.129P2-*Avil^tm2(cre)Fawa^*/J (Adv^Cre^)^132^, B6.129P2-*Nos2^tm1Lau^*/J (*Nos2*^-/-^)^133^, C57BL/6NJ-*Acod1^em1(IMPC)J^*/J (*Acod1*^-/-^), B6.129S-Tnftm1Gkl/J (*Tnf*^-/-^)^134^, B6.129S7-Ifngtm1Ts/J (*Ifng*^-/-^)^135^, and C57BL/6J-*^Spp1em1Msasn^*/J (*Spp1-IRES-TdTomato*)^136^ mice were purchased from Jackson Laboratories. B6-Fcgr1^tm2Ciphe^ (CD64^Cre^)^71^ mice were a gift from B. Malissen at Centre d’Immunologie de Marseille Luminy and were provided by Y. Belkaid at the National Institutes of Health. BM from *Il17*^-/-^*Il22*^-/-^ mice was a gift from S. Khader at the University of Chicago.

Il1rn^tm1.1Cga^ *(Il1rn*^fl^)^72^ were generated by Cem Gabay at the University of Geneva and were provided by A. Luster at Massachusetts General Hospital. *Sp140*^-/-^ mice were previously generated in-house^14^. *Sp140*^-/-^ *Ifnar1*^-/-^, *Acod1*^-/-^ *Ifnar1*^-/-^, and *Nos2*^-/-^ *Ifnar1*^-/-^ mice were generated by crossing *Sp140*^-/-^, *Acod1*^-/-^, and *Nos2*^-/-^ mice with *Ifnar1*^-/-^ mice in-house, respectively. *Sp140*^-/-^, *Acod1*^-/-^, and *Nos2*^-/-^ mice were crossed in-house with *Il1rn*^-/-^ mice to generate *Sp140*^-/-^ *Il1rn*^-/-^, *Acod1*^-/-^ *Il1rn*^-/-^, and *Nos2*^-/-^ *Il1rn*^-/-^ mice. *Sp140*^-/-^ *Il1rn*^fl/fl^ CD64^Cre^ mice were generated by crossing *Sp140*^-/-^ mice with *Il1rn*^fl/fl^ and CD64^Cre^ mice in-house. *Acod1*^-/-^ *Il1rn*^fl/fl^ Mrp8^Cre^ mice and *Nos2*^-/-^ *Il1rn*^fl/fl^ Mrp8^Cre^ mice were generated by crossing *Acod1*^-/-^ or *Nos2*^-/-^ mice with *Il1rn*^fl/fl^ and Mrp8^Cre^ mice in-house. Adv^Cre^ *Il1r1*^fl/fl^ and CD64^Cre^ *Il1r1*^fl/fl^ mice were generated by crossing Adv^Cre^ and CD64^Cre^ mice with *Il1r1*^fl/fl^ mice in-house.

### Bone marrow chimera mice

Adult mice were lethally irradiated with a Precision X-Rad320 X-Ray irradiator (North Branford, CT) at University of California, Berkeley and a gamma ray irradiator at WashU Medicine using a split dose of 500 rads each round with irradiation occurring 4-12 hours apart. Bone marrow (BM) from donor mice indicated in the figures was prepared by mashing femurs with a mortar and pestle to generate a single cell suspension. Red blood cells were then lysed with ACK lysis buffer (Thermo Fisher Scientific). The cells were then diluted in PBS and 1 × 10^7^ cells (either all one genotype or a 1: 1 mix of two genotypes for mixed BM chimera mice) were retro-orbitally injected into each recipient mouse. Mice were housed for at least 8 weeks post-injection to permit hematopoietic reconstitution prior to their use in experiments.

### *Mtb* infections

*Mtb* expressing Wasabi (*Mtb*-Wasabi) and *Mtb*-mCherry were generated using *Mtb* Erdman, a gift from S. A. Stanley, that had been passaged 2 or fewer times *in vitro*. *Mtb* was transformed with the pTEC15 plasmid (a gift from Lalita Ramakrishnan; Addgene plasmid # 30174)^137^ or the pMSP12::mCherry plasmid (a gift from Lalita Ramakrishnan; Addgene plasmid # 30167) to generate *Mtb*-Wasabi and *Mtb*-mCherry, respectively. For the transformation, *Mtb* was cultured in Middlebrook 7H9 liquid medium supplemented with 10% albumin-dextrose-saline, 0.4% glycerol, and 0.05% Tween-80 for 5 days at 37°C then washed in 10% glycerol to remove salts. Washed *Mtb* was electroporated with 1 µg DNA using a 2 mm electroporation cuvette and the following settings: 2500 volts, 1000 Ohms, 25 µF. Electroporated *Mtb* was grown on 7H11 plates supplemented with 10% oleic acid, albumin, dextrose, and catalase, 0.5% glycerol, and either 200 µg / mL Hygromycin for *Mtb*-Wasabi or 50 µg / mL Kanamycin for *Mtb*-mCherry for 3-4 weeks at 37°C. Colonies were picked and cultured in 10 mL inkwell flasks for 7 days at 37°C and then expanded into a 100 mL culture for 4-5 days at 37°C using 7H9 medium supplemented with 10% albumin-dextrose-saline, 0.4% glycerol, 0.05% Tween-80, and either Hygromycin for *Mtb*-Wasabi or Kanamycin for *Mtb*-mCherry. At log phase, the Mtb was filtered through a 5 µm syringe filter and frozen in 1 mL aliquots in 10% glycerol. Frozen cultures were diluted in distilled water to an O.D. of 0.0017 and then aerosolized with an inhalation exposure system (Glas-Col, Terre Haute, IN) thereby delivering ∼10-100 *Mtb* per mouse as determined by plating lungs day 1 post-infection to assess bacterial burden. The median day 1 counts across all experiments was 74.5 *Mtb* per mouse, with the quartiles being 32.75 and 117.

### Tissue processing for CFU and flow cytometry

Mice were harvested at 24-28 days following *Mtb* infection. Lungs were collected in gentleMACS C tube (Miltenyi Biotec) containing 3 mL of RPMI media with 70 µg / mL of Liberase TM (Roche) and 30 µg / mL of DNase I (Roche) and then processed into chunks using the lung_01 setting on a gentleMACS (Miltenyi Biotec).

Samples were then digested at 37°C for 30 minutes before being homogenized with the lung_02 setting on the gentleMACS. 2 mL of PBS with 20% Newborn Calf Serum (Thermo Fisher Scientific) were added to each sample to quench the digest before samples were filtered through 70 µm SmartStrainers (Miltenyi Biotec). Spleens were collected in 2 mL of PBS with 0.05% Tween-80 and then processed using mSpleen_01 setting on a gentleMACS (Miltenyi Biotec).

### Measuring bacterial burden

*Mtb* burden was assessed by serial dilution of the lung single cell suspension in phosphate-buffered saline (PBS) with 0.05% Tween-80. The dilutions were plated on 7H11 plates supplemented with 10% oleic acid, albumin, dextrose, and catalase and 0.5% glycerol and colonies were counted ∼3 weeks after plating.

### Flow cytometry of mouse tissue samples

Flow cytometry was performed by staining 100 µLs of lung single cell suspension with antibodies in PBS with 2% Newborn Calf Serum and 0.05% Sodium azide. A fixable viability dye (Ghost Dye Violet 510 or 540 or Red 780; Tonbo Biosciences), Super Bright Complete Staining Buffer (Thermo Fisher Scientific), and True-Stain Monocyte Blocker (BioLegend) were also added to every sample during surface antibody staining. Surface stains were incubated for 30-60 minutes at room temperature using a selection of the following antibodies: TruStain FcX PLUS (S17011E, BioLegend), BUV496-labeled CD45 (30-F11, BD Biosciences), BUV395-labeled CD11b (M1/70, BD Biosciences), BUV737-labeled CD11c (HL3, BD Biosciences), APC-R700-labeled Siglec F (E50-2440, BD Biosciences), PE-labeled MerTK (DS5MMER, Thermo Fisher Scientific), Super Bright 645-labeled MHC II (M5/114.15.2, Thermo Fisher Scientific), BV421-labeled PD-L1 (MIH5, BD Biosciences), BV711-labeled Ly6C (HK1.4, BioLegend), PE-Cy7-labeled MerTK (DS5MMER, Thermo Fisher Scientific), APC-labeled CD64 (X54-5/7.1, BioLegend), BUV563-labeled Ly6G (1A8, BD Biosciences), BV605-labeled MHC II (M5/114.15.2, BioLegend), Percp-Cy5.5-labeled CD9 (MZ3, BioLegend), eFluor 450-labeled Ly6D (49-H4, Thermo Fisher Scientific), BV480-labeled Fcer1a (MAR-1, BD Biosciences), BV650-labeled CD8*a* (53-6.7, BioLegend), BV785-labeled CD16.2 (9E9, BioLegend), AF594-labeled TREM2 (237920, R&D systems), AF647-labeled XCR1 (ZET, XCR1), APC-Fire 750-labeled CD19 (6D5, BioLegend), APC-Fire 750-labeled CD90.2 (30-H12, BioLegend), BUV615-labeled SIRP*a* (P84, BD Biosciences), BUV661-labeled B220 (RA3-6B2, BD Biosciences), PE-labeled Alcam (eBioALC48, Thermo Fisher Scientific), RB744-labeled CD9 (KMC8, BD Biosciences), BUV805-labeled PD-L1 (MIH5, BD Biosciences), and BUV805-labeled CD26 (H194-112, BD Biosciences). Following surface staining, samples were washed with PBS with 2% Newborn Calf Serum and 0.05% Sodium azide and then fixed for 20 minutes at room temperature with Cytofix/cytoperm (BD biosciences) prior to removal from the BSL3. If intracellular staining was performed, samples were incubated overnight at 4°C with combinations of the following antibodies: PE-Cy7-labeled CD63 (NVG-2, BioLegend), FITC-labeled C1q (JL-1, Thermo Fisher Scientific), AF700-labeled CD63 (NVG-2, BioLegend), eFluor 450-labeled Arg1 (A1exF5, Thermo Fisher Scientific), AF532-labeled iNOS (CXNFT, Thermo Fisher Scientific), and Percp-eFluor 710-labeled iNOS (CXNFT, Thermo Fisher Scientific). After washing out the intracellular stain, each sample received AccuCheck Counting Beads (Invitrogen) and was then analyzed on an Aurora (Cytek) flow cytometer. Data were analyzed with Flowjo version 10 (BD Biosciences).

### Confocal microscopy

A Zeiss LSM 880 laser scanning confocal microscope with 405, 458, 488, 514, 561, 594, and 633 lasers, with two photomultiplier detectors, a 34-channel GaASP spectral detector system, and a 2-channel AiryScan detector was used for microscopy. 20 µm sections were prepared from paraformaldehyde-fixed lungs from *Mtb*-mCherry-infected B6 and *Sp140*^-/-^ mice. The sections were stained overnight at 4°C with the following antibodies: BV421-labeled SIRP*a* (P84, BD Biosciences), Pacific Blue–labeled B220 (RA3-6B2, BioLegend), eF506-labeled CD4 (RM4-5, BioLegend), and AF647-labeled Ly6G (1A8, BioLegend). Diseased and healthy lesions were identified in stained sections with a 5× air objective and then imaged as a 20 µm z-stack acquired at a 1.5 µm step size with a 63× oil immersion objective lens with a numerical aperture of 1.4. The 63× oil immersion objective lens with a numerical aperture of 1.4 was also used to image single color-stained Ultracomp eBeads Plus (Thermo Fisher Scientific) for generating a compensation matrix.

### Image processing and histo-cytometry analysis

The Generate Compensations Matrix script^138^ was used in ImageJ to create a compensation matrix based on the single color-stained control images of the Ultracomp eBeads Plus (Thermo Fisher Scientific). The resulting compensation matrix was used for spectral unmixing of the lung images using Chrysalis software^138^, which was also used to rescale the data and generate new channels based on mathematical operations performed on existing channels. Imaris 9.9.1 (Bitplane) was used on the processed lung images to digitally identify cells based on protein expression using the surface creation tool and to measure the distance between *Mtb* and the nearest macrophage or neutrophil^139^. Statistics for the identified cells were exported from Imaris and then imported into FlowJo version 10 (BD Biosciences) for quantitative image analysis.

### Sorting Immune Cells for scRNA-seq analysis

For the myeloid dataset, lungs from 3 *Mtb*-Wasabi-infected as well as from 3 naïve *Acod1*^-/-^ and 3 naïve *Nos2*^-/-^ mice were processed as described for CFU and flow cytometry analysis 25 days after infection with the following slight modification: 3 minutes before harvesting, 0.25 µg of TotalSeq-A-labeled CD45.2 (104, BioLegend) was intravenously injected into each mouse to label cells in the vasculature^140^.

Surface staining of lung single cell suspensions was performed in PBS with 2% Newborn Calf Serum on ice for 30 minutes with the following antibodies: TruStain FcX PLUS (S17011E, BioLegend), APC-labeled Ly6G (1A8, BioLegend), APC-labeled CD64 (X54-5/7.1, BioLegend), and TotalSeq-A-labeled Ly6G (1A8, BioLegend). Magnetic enrichment was then performed with the EasySep APC Positive Selection Kit II (StemCell Technologies) and MojoSort Magnets (BioLegend) to increase the frequency of myeloid cells within the sample prior to sorting.

Following enrichment, samples were stained on ice for 45 minutes in PBS with 2% Newborn Calf Serum and True-Stain Monocyte Blocker (BioLegend) using a combination of the following antibodies: TotalSeq-A-labeled Ly6C (HK1.4, BioLegend), TotalSeq-A-labeled CD44 (IM7, BioLegend), TotalSeq-A-labeled CD274 (MIH6, BioLegend), TotalSeq-A-labeled Siglec F (S17007L, BioLegend), TotalSeq-A-labeled CSF1R (AFS98, BioLegend), TotalSeq-A-labeled CD11b (M1/70, BioLegend), TotalSeq-A-labeled CD86 (GL-1, BioLegend), TotalSeq-A-labeled MHC II (M5/114.15.2, BioLegend), TotalSeq-A-labeled CX3CR1 (SA011F11, BioLegend), TotalSeq-A-labeled CD11c (N418, BioLegend), TotalSeq-A-labeled CCR2 (SA203G11, BioLegend), TotalSeq-A-labeled CD62L (MEL-14, BioLegend), anti-mouse TotalSeq-A Hashtag antibody (1-6; BioLegend), PE-labeled Siglec F (S17007L, BioLegend), Pacific Blue-labeled B220 (RA3-6B2, BioLegend), and Pacific Blue-labeled CD90.2 (53-2.1, BioLegend). The TotalSeq-A-labeled antibodies detect protein expression in the scRNA-seq dataset, while the Hashtag antibodies allow several samples to be multiplexed together in a single lane on a 10X Genomics Chromium Next GEM Chip (cells from each mouse in a given 10X lane are labeled with a unique hashtag antibody allowing for deconvolution of mouse-specific responses during analysis)^141^. After staining, the samples were incubated with Sytox Blue Dead Cell Stain (Thermo Fisher Scientific) and sort purified using a 100 µm sorting chip in a 4 laser SH-800 cell sorter (Sony) on the purity setting. For *Mtb*-infected mice, *Mtb*-infected cells and bystander myeloid cells were individually purified, while samples from naïve lungs were sorted as macrophages and a mixture of neutrophils and monocytes that were then combined at a 1:2 ratio for greater representation of macrophages in the scRNA-seq dataset.

The stromal cell dataset was generated by infecting BM chimera mice made by transferring CD45.1 BM into lethally irradiated B6 or *Il1r1*^-/-^ hosts with *Mtb* 8 weeks post-reconstitution. Lungs from the infected animals as well as naïve chimera mice were harvested and processed as described for CFU and flow cytometry analysis. ACK lysing buffer (Thermo Fisher Scientific) was used to lyse the red blood cells in the samples, which were then stained for 30 minutes on ice with TruStain FcX PLUS (S17011E, BioLegend) and PE-Cy7-labeled CD45 (30-F11, BioLegend) antibody. The EasySep PE Positive Selection Kit II (StemCell Technologies) and MojoSort Magnets (BioLegend) were used to perform negative selection to enrich the CD45^-^ non-hematopoietic cells. The enriched cells were incubated with Sytox Green Dead Cell Stain (Thermo Fisher Scientific) and live CD45^-^ were sort purified using a 130 µm sorting chip in a 4 laser SH-800 cell sorter (Sony) on the purity setting.

The T cell and ILC datasets were generated by *Mtb* infection of mixed BM chimera mice generated by transferring WT CD45.1/2 and *Il1r1*^-/-^ BM into lethally irradiated CD45.1 mice.

Lungs were harvested 25 days post-infection and processed as described for measuring CFU and performing flow cytometry. Three mice were used for the T cell dataset, while the rarity of the ILCs required pooling lungs into 3 samples (2 samples containing 5 lungs and 1 sample containing 6 lungs). The T cell samples were treated with ACK lysing buffer (Thermo Fisher Scientific) to lyse the red blood cells and then stained on ice for 30 minutes with the following antibodies: BV421-labeled CD45.1 (A20, BioLegend), BV785-labeled CD45.2 (104, BioLegend), PE-labeled CD3e (17A2, BioLegend), APC-labeled CD11b (M1/70, Cytek), APC-labeled CD11c (N418, Cytek), APC-labeled CD19 (1D3, Cytek), TotalSeq-A-labeled Ly6C (HK1.4, BioLegend), TotalSeq-A-labeled CD44 (IM7, BioLegend), TotalSeq-A-labeled CD274 (MIH6, BioLegend), TotalSeq-A-labeled CX3CR1 (SA011F11, BioLegend), TotalSeq-A-labeled CD62L (MEL-14, BioLegend), TotalSeq-A-labeled TCRβ (H57-597, BioLegend), TotalSeq-A-labeled CD49a (, BioLegend), TotalSeq-A-labeled CD49a (HMα1, BioLegend), TotalSeq-A-labeled NKp46 (29A1.4, BioLegend), TotalSeq-A-labeled CD4 (RM4-5, BioLegend), TotalSeq-A-labeled ST2 (DIH9, BioLegend), TotalSeq-A-labeled CD25 (PC61, BioLegend), TotalSeq-A-labeled CD45.1 (A20, BioLegend), TotalSeq-A-labeled CD45.2 (104, BioLegend), TotalSeq-A-labeled CD8a (53-6.7, BioLegend), TotalSeq-A-labeled CXCR5 (L138D7, BioLegend), TotalSeq-A-labeled CCR6 (29-2L17, BioLegend), and anti-mouse TotalSeq-A Hashtag antibody (1-3; BioLegend). The stained T cell samples were then incubated with Sytox Green Dead Cell Stain (Thermo Fisher Scientific) and T cells (Live CD3e^+^ CD11b^-^ CD11c^-^ CD19^-^ CD45.1^+^ or CD45.2^+^ cells) were sort purified using a 100 µm sorting chip in a 4 laser SH-800 cell sorter (Sony) on the purity setting.

The ILC samples treated with ACK lysing buffer (Thermo Fisher Scientific) were stained on ice for 25 minutes with the following panel of lineage-specific antibodies: Biotin-labeled TER-119 (TER-119, BioLegend), Biotin-labeled GR-1 (RB6-8C5, BioLegend), Biotin-labeled CD3e (145-2C11, BioLegend), Biotin-labeled CD11b (M1/70, BioLegend), Biotin-labeled B220 (RA3-6B2, BioLegend), and Biotin-labeled TCRγ/δ (GL3, BioLegend). Lineage^-^ cells were enriched by negative selection using the EasySep Streptavidin RapidSpheres Isolation Kit (StemCell Technologies) and MojoSort Magnets (BioLegend). Enriched samples were then stained on ice for 30 minutes with the following antibodies: BV421-labeled CD45.1 (A20, BioLegend), BV785-labeled CD45.2 (104, BioLegend), PE-labeled CD127 (A7R34, BioLegend), Alexa Fluor 700-labeled CD127 (30-H12, BioLegend), Biotin-labeled CD5 (53-7.3, BioLegend), Biotin-labeled TCRβ (H57-597, BioLegend), TotalSeq-A-labeled Ly6C (HK1.4, BioLegend), TotalSeq-A-labeled CD44 (IM7, BioLegend), TotalSeq-A-labeled CD274 (MIH6, BioLegend), TotalSeq-A-labeled CX3CR1 (SA011F11, BioLegend), TotalSeq-A-labeled CD62L (MEL-14, BioLegend), TotalSeq-A-labeled TCRβ (H57-597, BioLegend), TotalSeq-A-labeled CD49a (, BioLegend), TotalSeq-A-labeled CD49a (HMα1, BioLegend), TotalSeq-A-labeled NKp46 (29A1.4, BioLegend), TotalSeq-A-labeled CD4 (RM4-5, BioLegend), TotalSeq-A-labeled ST2 (DIH9, BioLegend), TotalSeq-A-labeled CD25 (PC61, BioLegend), TotalSeq-A-labeled CD45.1 (A20, BioLegend), TotalSeq-A-labeled CD45.2 (104, BioLegend), TotalSeq-A-labeled CD8a (53-6.7, BioLegend), TotalSeq-A-labeled CXCR5 (L138D7, BioLegend), TotalSeq-A-labeled CCR6 (29-2L17, BioLegend), and anti-mouse TotalSeq-A Hashtag antibody (4-6; BioLegend). The enriched samples were then stained with APC-labeled Streptavidin (BioLegend) for 15 minutes on ice before being stained with Sytox Green Dead Cell Stain (Thermo Fisher Scientific). ILCs (Lineage^-^ IL-7Ra^+^ CD90.2^+^ CD45.1^+^ or CD45.2^+^ cells) were sort purified using a 100 µm sorting chip in a 4 laser SH-800 cell sorter (Sony) on the purity setting.

### Single cell RNA: Library generation and sequencing

The v3.1 chemistry Chromium Single Cell 3’ Reagent Kit (10X Genomics) was used to generate scRNA-seq libraries using the CITE-seq protocol^48^ modifications for the myeloid, T, and innate lymphoid cell libraries and the standard 10X protocol for the stromal cell library. The myeloid cell dataset required three lanes of a Chromium Next GEM Chip, with lane 1 containing naïve cells, lane 2 containing *Mtb*-infected cells, and lane 3 containing the bystander cells from the 3 *Acod1*^-/-^ and 3 *Nos2*^-/-^ mice. Each lane was super-loaded with ∼34,000 cells for a target of 17,000 single cells per lane at a multiplet rate of 3%^141^. The stromal cell dataset used 4 lanes of a Chromium Next GEM Chip, with lane 1 containing *Il1r1*^-/-^ stromal cells from *Mtb*-infected mice, lane 2 containing *Il1r1*^-/-^ stromal cells from naïve mice, lane 3 containing wild-type stromal cells from *Mtb*-infected mice, and lane 4 containing wild-type stromal cells from naïve mice. Each lane was loaded with 12,500 cells for an expected yield of 7,000 single cells at a multiplet rate of 5%. The T cell and ILC libraries were generated using the same Chromium Next GEM Chip, with lane 1 containing T cells and lane 2 containing ILCs from *Mtb*-infected mixed BM chimera mice. 25,000 cells were super-loaded into each lane with a target capture of 13,200 cells at a multiplet rate of 4%^141^. For the CITE-seq libraries, 0.5 U/µL RNaseOUT Recombinant Ribonuclease Inhibitor (Invitrogen) was added to single cell RT master mix during the loading step and 1 µL of ADT and HTO additive primers (0.2 µM stock) were added during the cDNA amplification. All samples were decontaminated by 2 rounds of centrifugation through 0.2 µM filter microcentrifuge tubes following cDNA, ADT, and HTO purification and then removed from the BSL3. The CITE-seq protocol was used to complete the ADT and HTO libraries while the 10X Genomics protocol was followed for the cDNA library. Quality control of the libraries was performed with a Fragment Analyzer (Agilent). Libraries were pooled and then sequenced on a NovaSeq 6000 (Illumina) using an S1 flow cell and the following cycles read 1 (28 cycles), i7 index (10 cycles), i5 index (10 cycles), read 2 (90 cycles).

### ScRNA-seq: data processing

Sequencing reads of the mRNA libraries were mapped to the mouse genome with CellRanger version 6.0.0 (10X Genomics), while the raw count matrices for the ADT and HTO libraries were generated with CITE-Seq-Count version 1.4.3^142^. The count matrices were analyzed with Seurat v4.3.0^47^ using default settings for normalizing the data (transcript data was log normalized while antibody reads were centered log-ratio transformed), finding variable features (the vst selection method was used to identify 2000 variable genes), and scaling the data. A detailed description of the settings used for the analysis is provided in the R code used for analysis (https://github.com/dmitrikotov/IL-1Ra-TB). For CITE-seq datasets, multiplets were excluded through HTO demultiplexing using the HTODemux function (positive quantile set at 0.99, above which a cell is defined as positive). The myeloid cell dataset was filtered to include cells with between 200 and 4500 genes and less than 5% mitochondrial reads. The T cell, ILC, and stromal cell datasets were filtered to only include cells that have between 200 and 2500 genes and less than 5% mitochondrial reads. All dataset integrations were performed using anchor-based CCA integration. The resulting myeloid dataset as well as our previously published scRNA-seq datasets for myeloid cells in *Mtb*-infected and naïve B6 and *Sp140*^-/-^ mice (GSE216023) were integrated together and then clustered with a resolution of 1.5. The T cell and ILC datasets were analyzed with the default Seurat workflow using weighted nearest neighbor multimodal analysis to cluster cells based on mRNA and protein expression at a resolution of 1. The stromal cell dataset was also analyzed with Seurat using integration to combine the *Mtb*-infected and naïve WT and *Il1r1*^-/-^ datasets and then clustering the cells based on mRNA expression using a resolution of 1.5.

### Bulk RNA-seq, Gene signature, and ROC curve analysis

Our previously published signatures for type I IFN and IFN-γ^15^ were applied to our myeloid cell scRNA-seq dataset to score cells based on their gene expression using the UCell package^143^. Signatures for the B6, *Nos2*^-/-^, *Acod1*^-/-^, and *Sp140*^-/-^ mice were generated in R^144^ by identifying genes induced at least 8-fold in the myeloid cells from *Mtb*-infected mice relative to naïve animals for each genotype. A corresponding human signature was produced using the convert_mouse_to_human_symbols from the NicheNetr package^76^. The hacksig package using the singscore statistical method^148^ was applied to calculate a signature score based on each signature’s expression in the TANDEM cohort PBMC RNA-seq data^49^, Berry South Africa RNA-seq data^50^, or the Bloom et al. microarray data^51^. The pROC package^146^ was used to generate the ROC curves and calculate the area under the curve to quantify the ability of each signature to discriminate between the active *Mtb* cases and healthy controls in the TANDEM cohort, between active *Mtb* cases and LTBI individuals in the Berry South Africa dataset, and between active *Mtb* cases and lung cancer or pneumonia in the Bloom et al. dataset.

We also used the caret^147^ package in R for generating additional signatures for each mouse genotype, using machine learning feature selection with the input being differentially expressed genes (DEGs) defined as genes induced at least 2-fold and with an adjusted p-value below 0.05 in *Mtb*-infected relative to naïve lungs (B6 = 1071, *Sp140*^-/-^ = 585, *Acod1*^-/-^ = 474, *Nos2*^-/-^ = 502 DEGs). The NicheNetr package’s convert_mouse_to_human_symbols function was used to convert the mouse DEGs to human symbols. Using caret, recursive feature elimination was performed on the DEGs to identify potential features (B6 = *GBP2*, *CASP1*, *BATF2*, *CASP4*, *SCARF1*, *CD274*, *NMI*, *GBP6*, *GBP5*, *RUNX3*, *GADD45B*, *PDCD1LG2*, *MS4A6A*, *NPC2*, *UBE2L6*, *SECTM1*, *TAP1*, *IRF1*, *BEX3*, *AGTRAP*, *C1QA*, *IFI30*, *C1QC*, *B2M*, *TIMM10*; *Sp140*^-/-^ = *GBP2*, *CASP1*, *BATF2*, *CASP4*, *SCARF1*, *CD274*, *MAP4*, *GBP6*, *SOCS3*, *RAB20*, *MS4A6A*, *GBP5*, *FBXO6*, *NMI*, *PDCD1LG2*, *NPC2*, *TAP1*, *UBE2L6*, *GADD45B*, *IRF1*; *Acod1*^-/-^ = *GBP2*, *CASP1*, *CASP4*, *BATF2*, *SCARF1*, *CD274*, *PDCD1LG2*, *GBP6*, *NMI*, *NPC2*, *MS4A6A*, *SOCS3*, *GBP5*, *B2M*, *IRF1*, *TAP1*, *AGTRAP*, *SECTM1*, *HK3*, *UBE2L6*; *Nos2*^-/-^ = *GBP2*, *CASP1*, *CASP4*, *BATF2*, *SCARF1*) for building a classification model using a Random Forest algorithm with the TANDEM cohort as the training dataset. The final models for each genotype were tested on the Berry South Africa cohort, with the pROC package used to generate the ROC curves and calculate the area under the curve to quantify the ability of each signature to discriminate between active *Mtb* cases and LTBI individuals in the Berry South Africa dataset.

The hacksig and pROC packages were also used to test the predictive ability of a gene signature consisting of known immunosuppressive genes (*IL1RN*, *IL18BP*, *CD274*, *TGFB1*, *IDO1*, *IL10*, and *ARG1*) on the TANDEM cohort. The pROC package was also used to determine the ability of *IL1RN* expression to identify active *Mtb* cases in the TANDEM cohort and the Berry South Africa dataset. The DESeq2 package^148^ was used to analyze *IL1RN* expression in human PBMCs from *Mtb*-infected or healthy control individuals in the TANDEM cohort as well as to analyze *IL1RN* expression relative to time prior to TB diagnosis in the progressor portion of the Adolescent Cohort Study^61^. The pROC package was also used to compare the relative performance of different gene signatures.

### NicheNet analysis of IL-1-regulated signaling

IL-1-regulated genes were defined as genes expressed at least 2-fold higher in WT than *Il1r1*^-/-^ cells in the T cell, innate lymphoid cell, and stromal cell scRNA-seq datasets. These IL-1-regulated genes were used as the potential ligands for NicheNet analysis^76^. The receiver cells were defined as IMs, *Trem2*^+^ IMs, and *Spp1*^+^ IMs from the myeloid cell scRNA-seq dataset. The expressed receptors on the receiver cells were identified using the get_expressed_genes function from NicheNet with a 10% cutoff for expression and then filtering the NicheNet receptor database to those genes expressed in the IM populations. Ligand activity in the receiver cells was calculated by identifying genes upregulated in *Trem2*^+^ or *Spp1*^+^ IMs relative to the IM cell population via the FindMarkers function from the Seurat package^47^, focusing on genes upregulated at least 2-fold and with an adjusted p value less than 0.05, as calculated by the Wilcoxon Rank-Sum test with Bonferroni correction. The ability of each IL-1-regulated ligand to induce the genes upregulated in *Trem2*^+^ or *Spp1*^+^ IMs was determined by NicheNet’s predict_ligand_activities function.

### *In vitro* bone marrow-derived macrophage (BMDM) differentiation

Murine BM cells were isolated from femurs and tibias by crushing the bones with a mortar and pestle in complete DMEM (DMEM (Gibco) supplemented with 10% heat-inactivated Fetal Bovine Serum (Seradigm, Avantor), 2% HEPES, 1mM sodium pyruvate, 100U/mL penicillin/streptomycin, and 2mM GlutaMAX (Gibco)). Cell suspensions were filtered through a 70µM filter (Corning), and red blood cells were lysed with ACK lysis buffer (Thermo Fisher Scientific). BM cells were differentiated in complete DMEM for 4 days with 40 ng/µL of M-CSF (Peprotech) at 37°C and 5% CO^2^. After 4 days, cells were harvested and 2.5 x 10^5^ cells were plated per well in a 6-well plate. Differentiation was continued with 20 ng/µl M-CSF, 10 ng/µL IL-4 (Peprotech), 10 ng/uL TGF-β (Sino Biological), and / or 40 ng/µL GM-CSF (Peprotech) for 3 days.

### Flow cytometry of *in vitro* cultures

Differentiated BMDMs were harvested and resuspended in 100µl of PBS with 2% Newborn Calf Serum and 0.05% Sodium azide for surface staining.

Samples were incubated with RB670-labeled MerTK (BB14-16, BD Biosciences), BV650-labeled CD64 (X54-5/7.1, BD Biosciences), BUV737-labeled CD11c (HL3, BD Biosciences), BV711-labeled PD-L1 (MIH5, BD Biosciences), and RY703-labeled CD9 (KMC8, BD Bioscience) antibodies as well as TruStain FcX PLUS (S17011E, BioLegend), Ghost Dye 780 (Cytek Biosciences) and Brilliant Stain Buffer Plus (BD, Biosciences) for 1 hour at room temperature. Following surface staining, samples were washed with PBS with 2% Newborn Calf Serum and 0.05% Sodium azide, fixed for 20 minutes at room temperature with 4% paraformaldehyde, then washed and resuspended with PermWash (BD Biosciences). For intracellular staining, samples were incubated with PE-Cy7-labeled CD63 (NVG-2, BioLegend) overnight at 4°C.

### Quantification of IL-1Ra secretion

Supernatant from *in vitro* differentiated BMDMs was collected and stored at-20°C for no longer than two weeks. Samples were not subject to multiple freeze-thaw cycles. IL-1Ra protein was quantified using the mouse IL-1ra/IL-1F3 Quantikine enzyme-linked immunosorbent assay (ELISA) kit (MRA00, R&D Systems) according to the manufacturer’s instructions.

### Statistical analysis

Statistical significance was determined using Prism (GraphPad) software for unpaired or paired two-tailed Student t test when comparing two populations, one-way or two-way ANOVA tests with Tukey’s multiple comparisons test when comparing multiple groups, and by Pearson’s correlation coefficient when examining the correlation of two parameters.

ROC curves, their confidence intervals, and their AUC were calculated in R using the pROC package.

## Data availability

Raw and processed single cell RNA-sequencing data is deposited at NCBI Gene Expression Omnibus: GSE255213, GSE254983, and GSE254926.

## Code availability

Code for scRNA-sequencing analysis is available on Github: https://github.com/dmitrikotov/IL-1Ra-TB.

## Extended Data Figure Legends

**Extended Data Figure 1. Neutrophils and interstitial macrophages localize to diseased tissue, with macrophages localizing closer to *Mtb*.** (**A**) Number and (**B**) frequency among total lung cells of neutrophils, monocytes, IMs, and AMs in the lungs of *Mtb*-infected *Sp140*^-/-^ (n = 13), *Sp140*^-/-^ *Ifnar1*^-/-^ (n = 11), *Acod1*^+/+^ (n = 7), *Acod1*^+/-^ (n = 6), *Acod1*^-/-^ (n = 12), *Nos2*^+/+^ (n = 4), *Nos2*^+/-^ (n = 7), and *Nos2*^-/-^ (n = 8) mice. (**C**) Representative images and (**D**) image quantification of *Mtb* (*Mtb*-mCherry^+^, red), macrophages (SIRPα^+^, blue), and neutrophils (Ly6G^+^, green) in diseased and healthy portions of *Mtb*-infected B6 (n = 5) and *Sp140*^-/-^ lungs (n = 6). (E) Representative images and (**F**) image quantification of *Mtb* (*Mtb*-mCherry^+^, purple), macrophages (SIRPα^+^, green), CD4^+^ T cells (CD4^+^, yellow), and neutrophils (Ly6G^+^, red) in diseased and healthy portions of *Mtb*-infected B6 (n = 6), *Acod1*^-/-^ (n = 6), and *Nos2*^-/-^ lungs (n = 6). (**G**) Quantification of the distance between macrophages or neutrophils to the nearest *Mtb* in representative images of B6, *Sp140*^-/-^, *Acod1*^-/-^, and *Nos2*^-/-^ lungs. Lungs were analyzed for the experiments depicted in (A) – (G) 24-26 days after *Mtb*-Wasabi or *Mtb*-mCherry infection. The bars in (A) and (B) represent the median. Pooled data from two independent experiments are shown in (A), (B), (D), and (F). Statistical significance in (A) and (B) by one-way ANOVA with Tukey’s multiple comparison test, in (D) and (F) by paired t test, and in (G) by two tailed t test. *p < 0.05, **p < 0.01, ***p < 0.001, ****p < 0.0001.

**Extended Data Figure 2. Classifying myeloid scRNA-seq clusters and comparing myeloid cells from naïve *Sp140*-, *Nos2*-, and *Acod1*-deficient mouse lungs. Dot plot of lineage of markers used to** (**A**) identify myeloid cell clusters and (**B**) define neutrophil cluster heterogeneity. (**C**) The myeloid populations in the naïve lungs of B6, *Nos2*^-/-^, *Acod1*^-/-^, and *Sp140*^-/-^ mice (n = 3 per genotype). (**D**) Number of shared and unique genes differentially expressed (2-fold change and adjusted p value < 0.05) between by neutrophils, alveolar macrophages, monocytes, and interstitial macrophages in *Nos2*^-/-^, *Acod1*^-/-^, and *Sp140*^-/-^ relative to those from B6 mice. Statistical significance in (D) was calculated with the Wilcoxon Rank-Sum test with Bonferroni correction.

**Extended Data Figure 3. Mouse gene signatures consisting of genes upregulated in myeloid cells following *Mtb* infection of B6, *Sp140*^-/-^, *Acod1*^-/-^, or *Nos2*^-/-^ mice discriminate active *Mtb* infection from Cancer and Pneumonia in humans.** (**A**) A heatmap depicting the log_2_ fold change of gene upregulation following *Mtb* infection relative to the gene’s expression in myeloid cells from naïve lungs. The heatmap compares gene expression for signature genes between B6, *Nos2*^-/-^, *Acod1*^-/-^, and *Sp140*^-/-^ mice (n = 3 per genotype). Grey boxes indicate genes not present in the gene signature of a given mouse genotype. Genes in blue indicate those with known immunosuppressive function. (**B**) ROC curves of the ability of machine learning-generated signatures consisting of genes induced by *Mtb* infection in B6, *Nos2*^-/-^, *Acod1*^-/-^, or *Sp140*^-/-^ mice to stratify human active *Mtb* patients (n = 16) and LTBI individuals (n = 31) in the Berry South Africa cohort (analysis of GSE107992). (**C**) ROC curves of the ability of genes induced by *Mtb* infection in B6, *Nos2*^-/-^, *Acod1*^-/-^, or *Sp140*^-/-^ mice to stratify human active *Mtb* patients (n = 35) and individuals with lung cancer (n = 16) or Pneumonia (n = 19) in the Bloom et al. cohort (analysis of GSE42834). Parenthetical numbers indicate the 95% confidence interval, and p-values are for comparisons between the given susceptible mouse signature and the B6 signature for (B) and (C). Statistical significance in (B) and (C) was calculated with DeLong’s test.

**Extended Data Figure 4. Classifying scRNA-seq clusters for *Mtb*-infected non-human primate and human lung datasets and utilizing the human dataset to identify a gene signature conserved between mouse and human *Spp1*^+^ IMs.** Dot plot of lineage of markers used to identify cell clusters in (**A**) *Mtb*-infected non-human primate and (**B**) human lungs. (**C**) Venn diagram depicting the number of genes preferentially expressed (2-fold change and adjusted p value < 0.05) by *Spp1*^+^ IMs relative to *Trem2*^+^ IMs in *Mtb*-infected B6, *Nos2*^-/-^, *Acod1*^-/-^, and *Sp140*^-/-^ mouse lungs (n = 3 per genotype) and human lungs (analysis of GSE192483). (**D**) Core gene signature for genes commonly expressed by *Spp1*^+^ IMs in all three susceptible mouse models as well as those expressed by the three susceptible mouse models and humans.

**Extended Data Figure 5. IL-1Ra expression in mice and humans.** (**A**) Myeloid cell expression of *Il1r2* and *Il1rn* in naïve and *Mtb*-infected lungs from B6, *Nos2*^-/-^, *Acod1*^-/-^, and *Sp140*^-/-^ mice (n = 3 mice per genotype for naïve lungs and 3 mice per genotype for *Mtb*-infected lungs) as measured by scRNA-seq. (**B**) *Il1rn* expression in myeloid cells from the lungs of B6, *Nos2*^-/-^, *Acod1*^-/-^, and *Sp140*^-/-^ mice (n = 3 mice per genotype for naïve lungs and 3 mice per genotype for *Mtb*-infected lungs). (**C**) Normalized *IL1RN* expression in *Mtb*-infected (n = 151) and healthy individuals (n = 98) from the TANDEM cohort (analysis of GSE114192). (**D**) ROC curve of the predictive ability of an immunosuppressive gene signature applied to the TANDEM cohort. Parenthetical numbers indicate the 95% confidence interval. (**E**) *IL1RN* expression plotted relative to time before clinical diagnosis of *Mtb* infection in the progressor population of the Adolescent Cohort Study (n = 74) with the blue line indicating local polynomial regression fitting using the loess method and grey indicating the 95% confidence interval. Day 200 prior to diagnosis (red line) indicates the timepoint *IL1RN* expression started to increase in patients as they approached diagnosis (analysis of GSE79362). Statistical significance in (A) was calculated with the Wilcoxon Rank-Sum test with Bonferroni correction, and in (C) was calculated using the Wald test and corrected for multiple testing using the Benjamini and Hochberg method. ****p < 0.0001.

**Extended Data Figure 6. Flow cytometric identification of monocyte and monocyte-derived macrophage subsets identified by scRNA-seq.** (**A**) Overview of monocyte and interstitial macrophage subsets and (**B**) their expression of subset identifying markers as measured by scRNA-seq of *Mtb*-infected lungs from B6, *Nos2*^-/-^, *Acod1*^-/-^, and *Sp140*^-/-^ mice (n = 3 mice per genotype). (**C**) Myeloid cell flow cytometry gating strategy. (**D**) Frequency of *Mtb*-infected cells among the major myeloid cell populations. (**E**) Gating of mature and immature IMs defined by expression of CD11c and PD-L1 and (**F**) flow cytometric analysis of CD64, Ly6C, PD-L1, and MHC II expression by immature (n = 27) and mature (n = 27) IMs in *Mtb*-infected lungs (day 27 after infection). The bars in (F) represent the median. Pooled data from two independent experiments are shown in (F). Statistical significance in (F) was calculated by two-tailed t test. ****p < 0.0001.

**Extended Data Figure 7. The impact of mouse genotype and timing on monocyte-derived macrophage subsets in *Mtb*-infected lung.** (**A**) Quantification of the number of the total mature IMs, (**B**) mature *Spp1*^+^ IMs, and (**C**) mature *Trem2*^+^ IMs in the lungs of *Mtb*-infected B6 (n = 9), *Sp140*^-/-^ (n = 16), *Nos2*^-/-^ (n = 13), and *Acod1*^-/-^ (n = 9) mice. (**D**) scRNA-seq analysis of C3H mouse lung parenchymal monocytes and monocyte-derived macrophages at days 0, 10, 17, and 34 after *Mtb* infection (analysis of GSE265843). (**E**) Flow cytometric analysis of Wasabi gMFI in *Mtb*-Wasabi harboring total (immature and mature) IM, *Spp1*^+^ IM, and *Trem*2^+^ IM in the lungs of B6 (n = 13) and *Sp140*^-/-^ mice (n = 12). (**F**) scRNA-seq analysis of Nos2 and Arg1 expression in monocytes and macrophages from *Mtb*-infected lungs of B6, *Sp140*^-/-^, *Nos2*^-/-^, and *Acod1*^-/-^ mice (n = 3 per genotype). (**G**) Representative histogram of Arg1 and iNOS expression by mature *Trem2*^+^ IM (blue), mature *Spp1*^+^ IM (red), mature IM (green), and monocytes (grey) from *Mtb*-infected mouse lungs. For (A) – (C) and (E), lungs were analyzed 24-28 days after *Mtb* or *Mtb*-Wasabi infection. The bars in (A) – (C) and (E) represent the median. Pooled data from two independent experiments are shown in (A) – (C) and (E). Statistical significance in (A) – (C) was calculated by two-tailed t test and in (H) by one-way ANOVA with Tukey’s multiple comparison test. *p < 0.05, **p < 0.01, ***p < 0.001.

**Extended Data Figure 8. Myeloid cells express very little IL-1 receptor and IM as well as neuronal expression of IL-1 receptor does not impact bacterial control in a cell-intrinsic manner.** (**A**) The myeloid populations identified by scRNA-seq and (**B**) *Il1r1* expression in myeloid cells from the lungs of B6, *Nos2*^-/-^, *Acod1*^-/-^, and *Sp140*^-/-^ mice (n = 3 mice per genotype for naïve lungs and 3 mice per genotype for *Mtb*-infected lungs). (**C**) Mixed BM chimera schematic, (**D**) representative flow plot, and (**E**) quantification of the frequency of *Mtb* infection among lung wild-type CD45.1^+^ and *Il1r1*^-/-^ CD45.1^-^ IMs (n = 8) 25 days post-infection. (F) Experimental setup and (**G**) lung bacterial burden of bone marrow chimera mice with wild-type neurons (CD45.1 host, n = 13), littermate control neurons (Adv^+/+^ Il1r1^fl/fl^ host, n = 9), and IL-1 receptor-deficient neurons (Adv^Cre/+^ Il1r1^fl/fl^ host, n = 14) 25 days after *Mtb* infection. Pooled data from two independent experiments is shown in (E) and (G). Statistical significance in (E) was calculated with a paired t test and in (G) with a one-way ANOVA with Tukey’s multiple comparison test.

**Extended Data Figure 9. Classifying T cell, ILC, and stromal cell scRNA-seq clusters.** Dot plot of lineage of markers used to annotate the clusters in our (**A**) T cell dataset, (**B**) ILC dataset, and (**C**) stromal cell dataset.

**Extended Data Figure 10. NicheNet analysis of IL-1 regulated ligands that could activate interstitial macrophages.** (**A**) List of IL-1 regulated potential ligands identified in stromal and hematopoietic cells. (**B**) Receptor expression on interstitial macrophages for the IL-1-regulated ligands. (**C**) Predicted ligand activity based on genes differently expressed by the activated interstitial macrophage subsets. (**D**) Quantification of the total number of innate immune cells (neutrophils, monocytes, IMs, and AMs) and (**E**) adaptive immune cells (B cells, CD4^+^ T cells, and CD8^+^ T cells) in the lungs of *Mtb*-infected mixed BM chimera mice that received *Tnf*^-/-^: CD45.1 mixed BM (n = 7), *Il1r1*^-/-^: CD45.1 mixed BM (n = 9), *Tnf*^-/-^: *Il1r1*^-/-^ mixed BM (n = 18), and *Il1r1*^-/-^: *Il1r1*^-/-^ BM (n = 14). Lungs were analyzed 25-26 days after *Mtb* or *Mtb*-Wasabi infection. The bars in (D) and (E) represent the median. Pooled data from two independent experiments are shown in (D) and (E). Statistical significance in (D) and (E) was calculated with a two-way ANOVA with Geisser-Greenhouse correction and Tukey’s multiple comparison test. *p < 0.05, **p < 0.01, ***p < 0.001, ****p < 0.0001.

