## Supplementary Figures for "Interleukin-1 receptor antagonist is a conserved factor for exacerbating tuberculosis susceptibility"

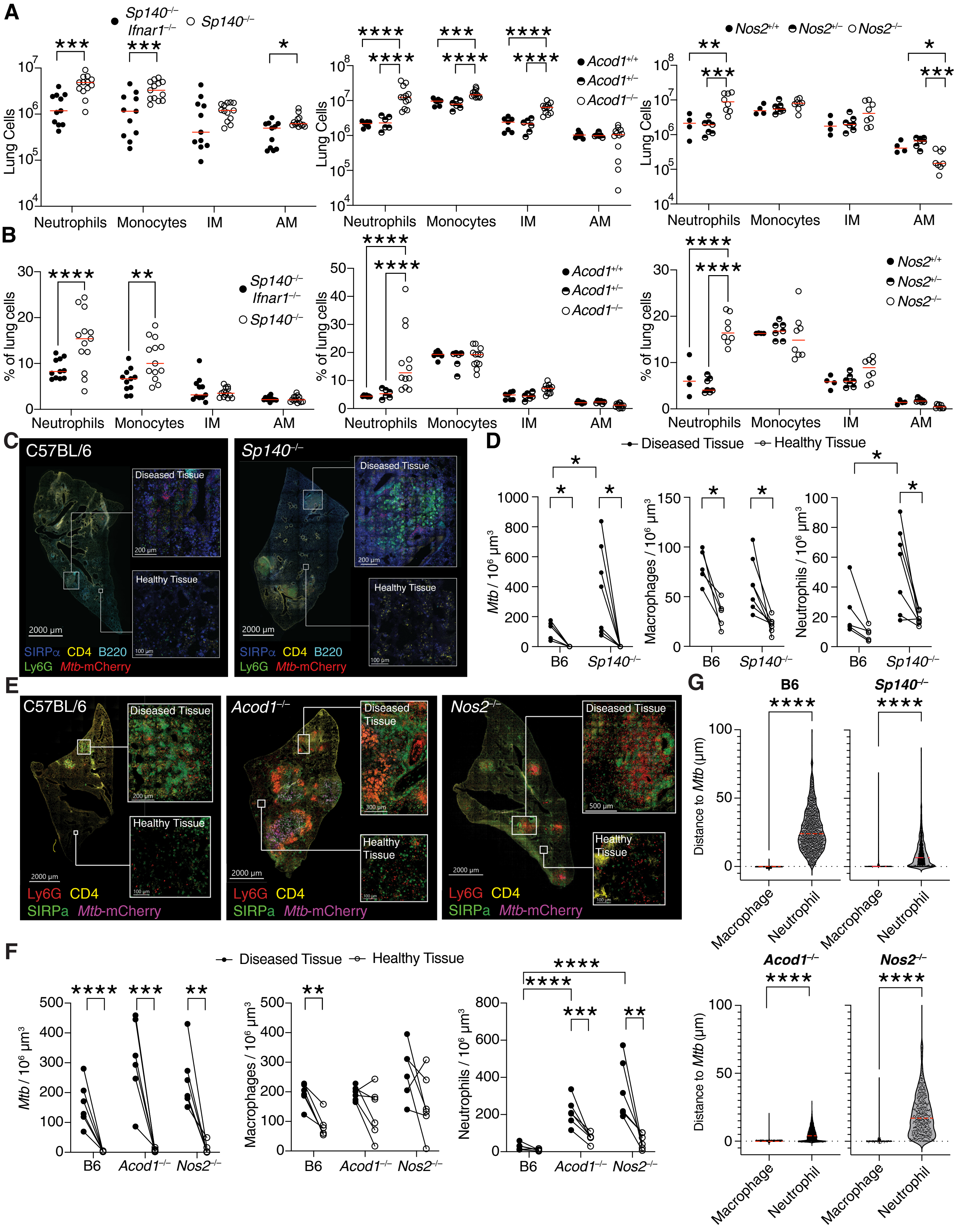


**Extended Data Figure 1.**

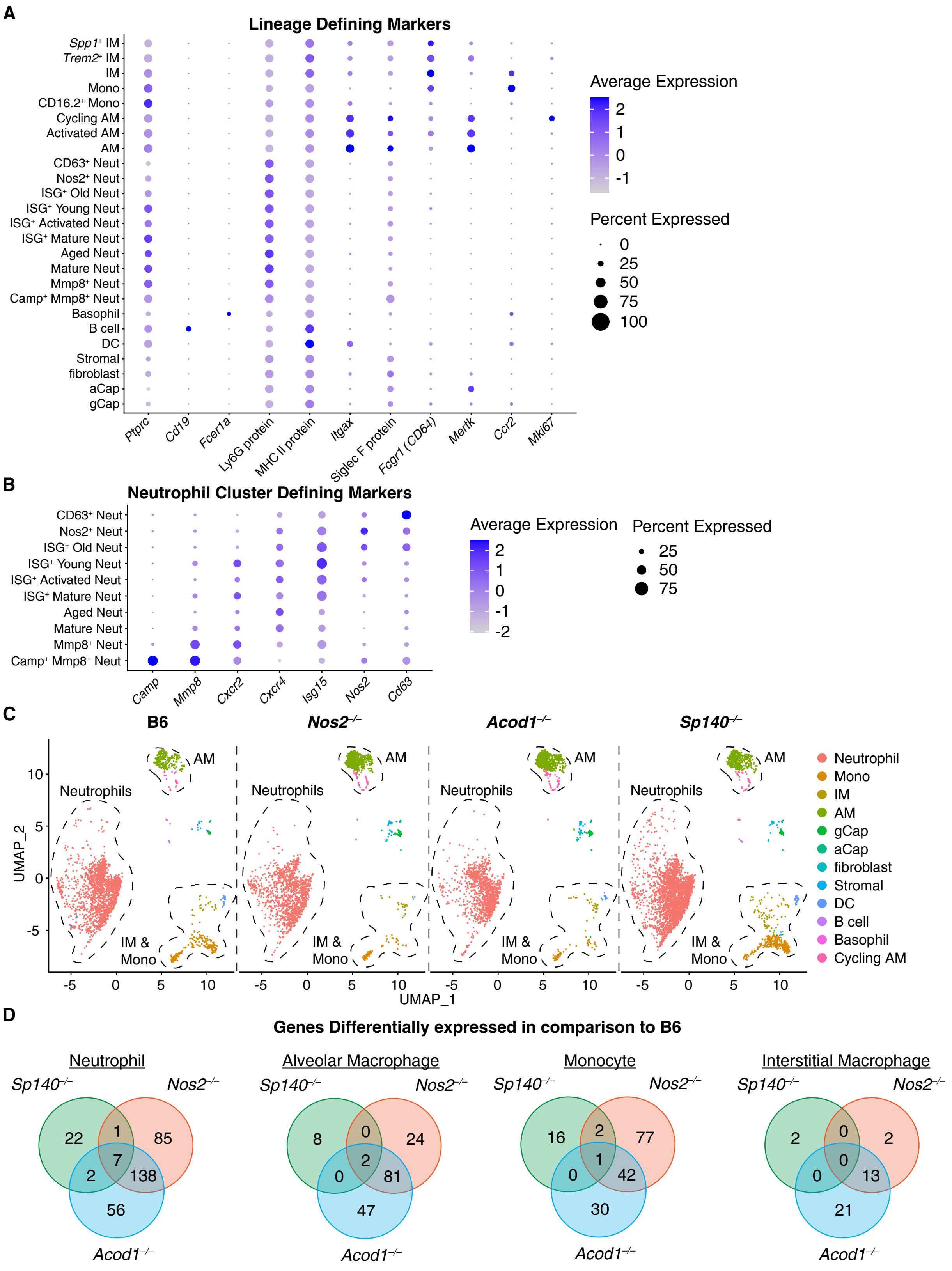


**Extended Data Figure 2.**

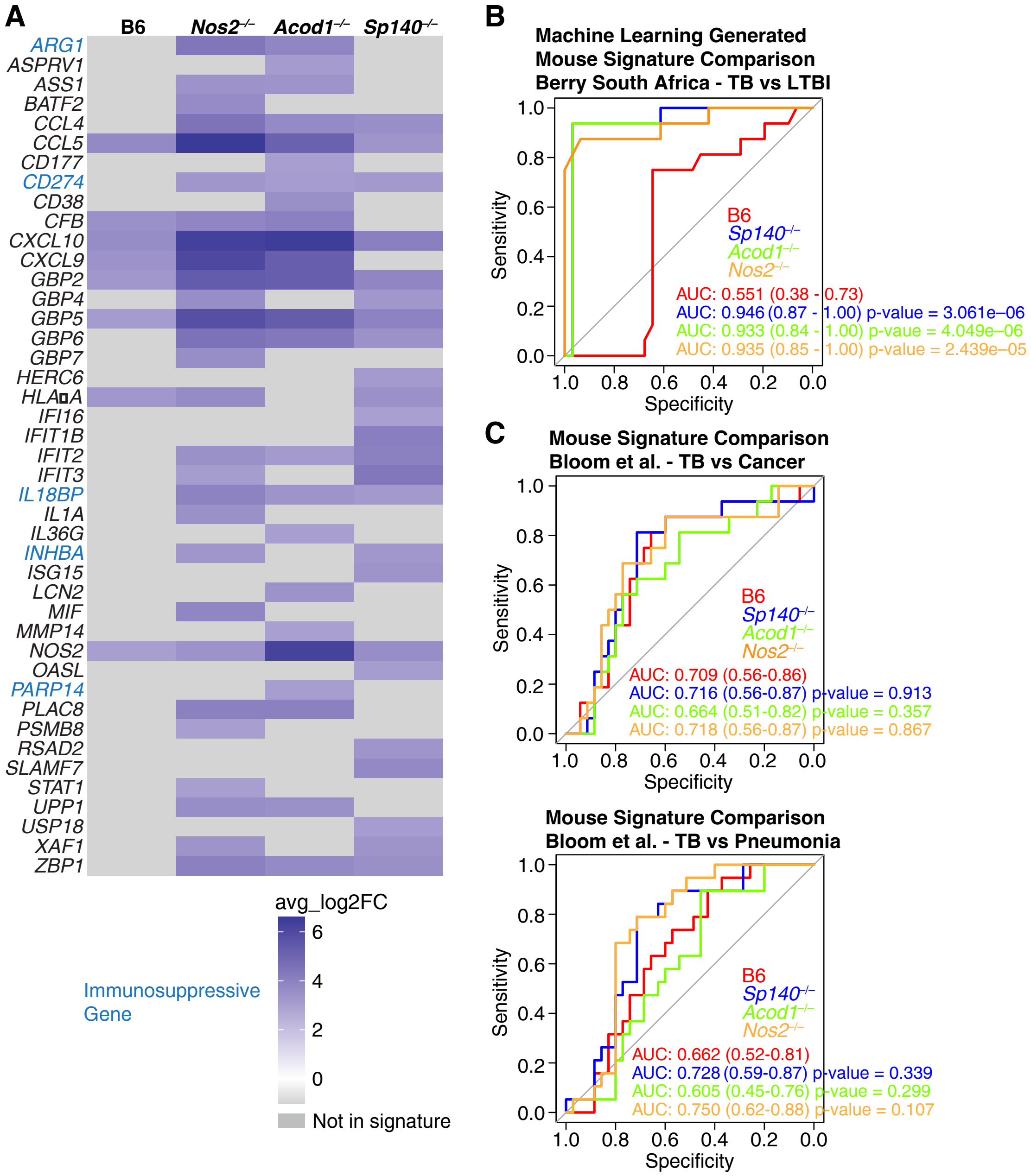


**Extended Data Figure 3.**

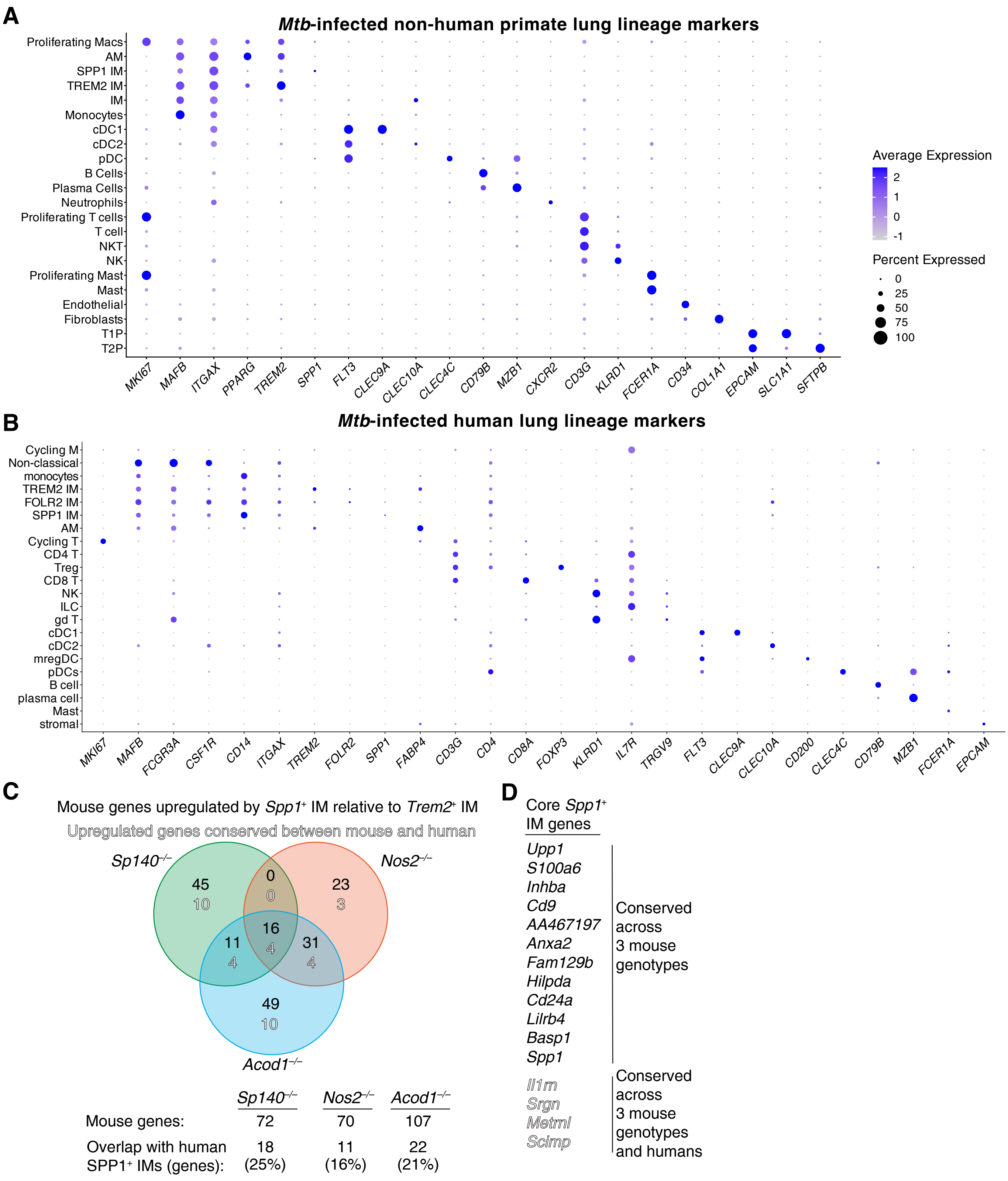


**Extended Data Figure 4.**

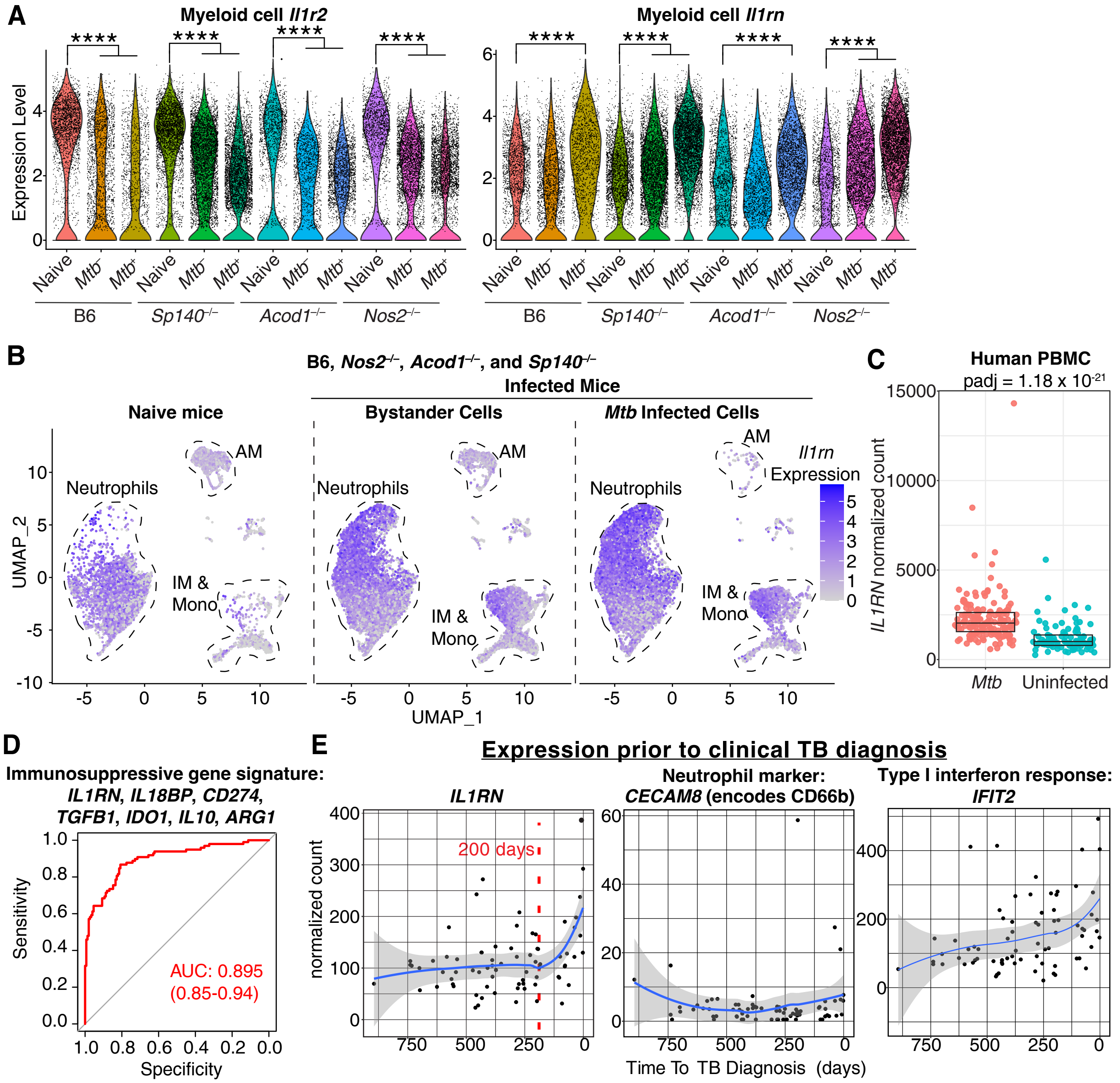


Extended Data Figure 5.

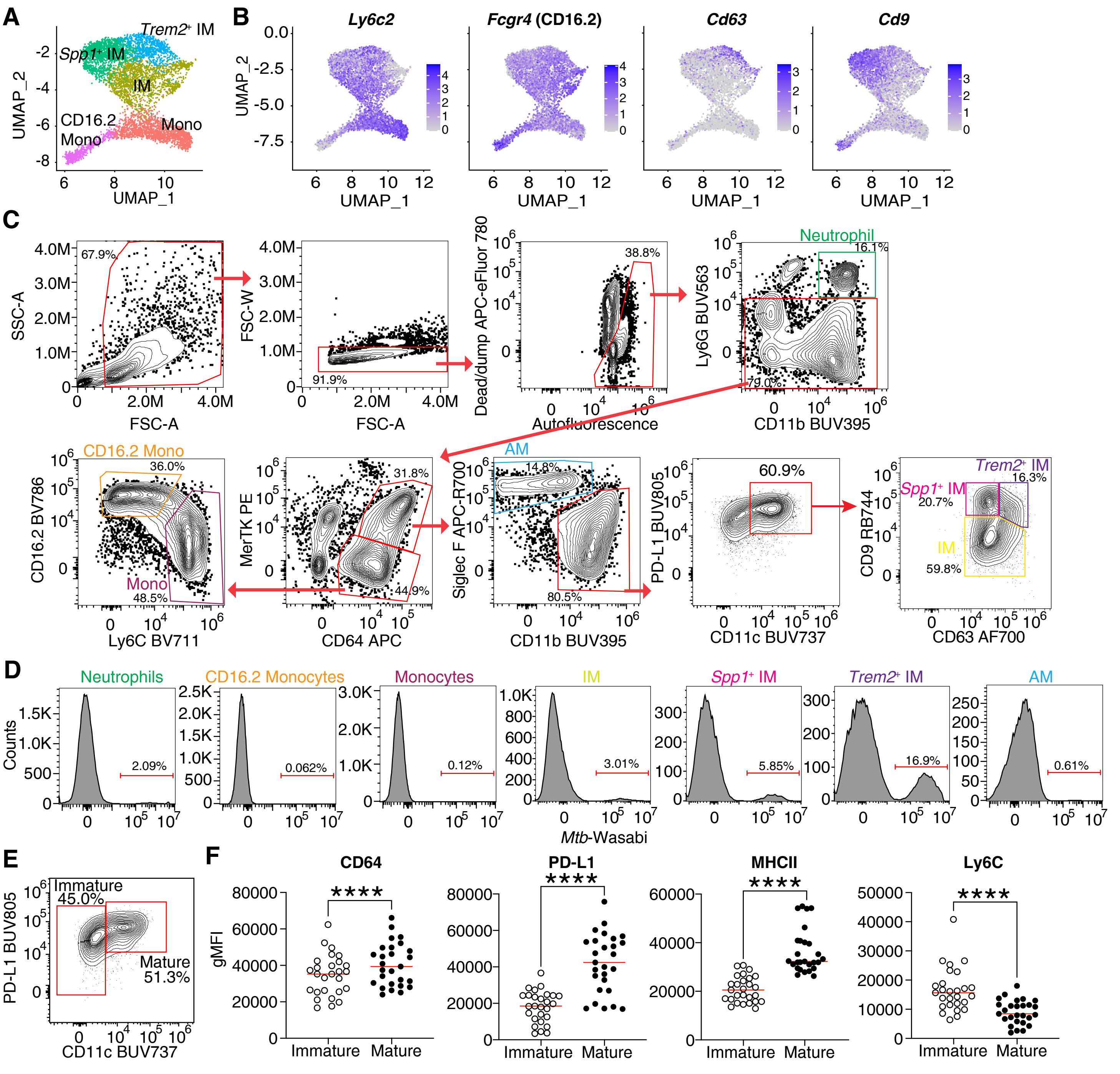


**Extended Data Figure 6.**

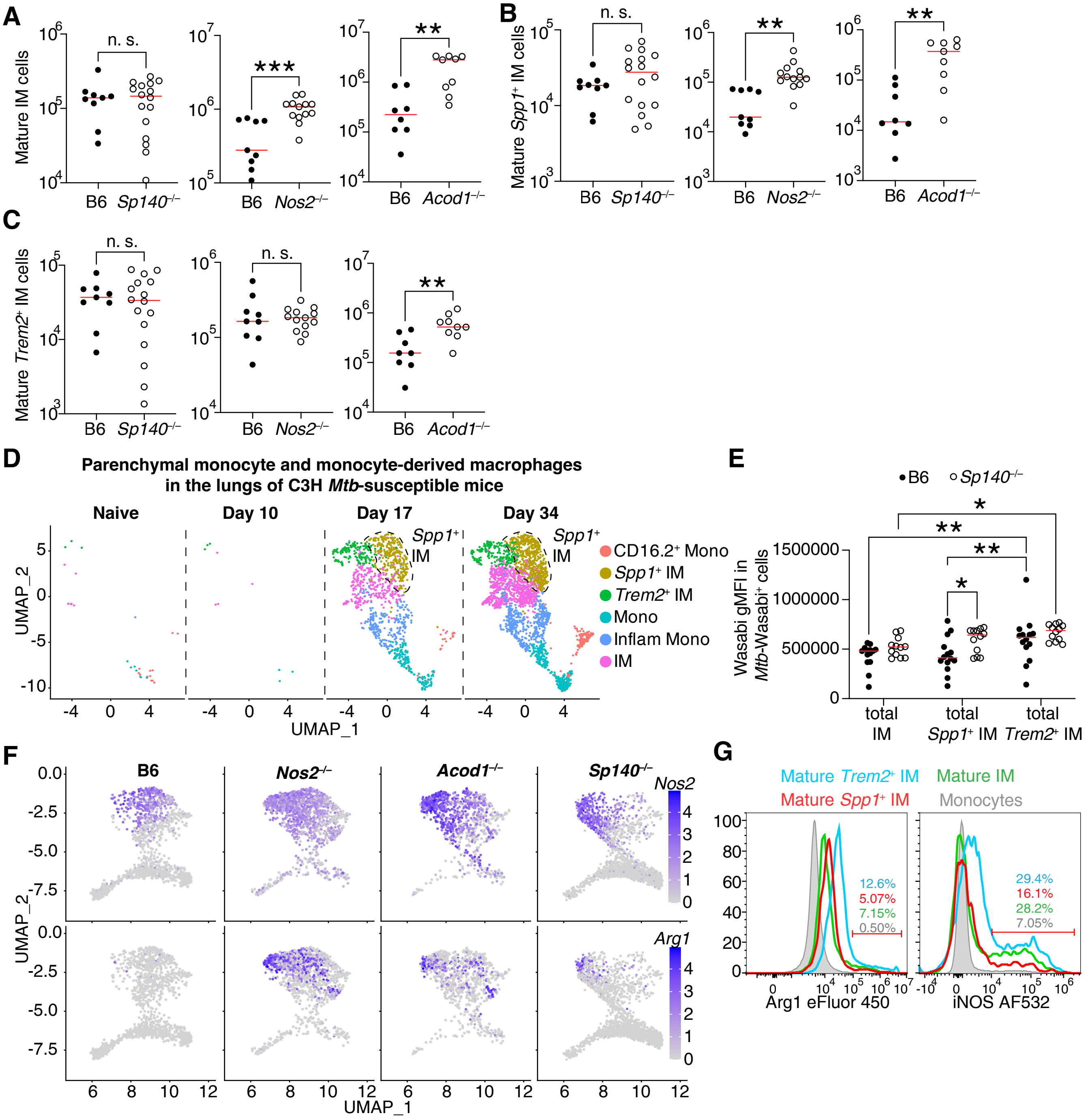


Extended Data Figure 7.

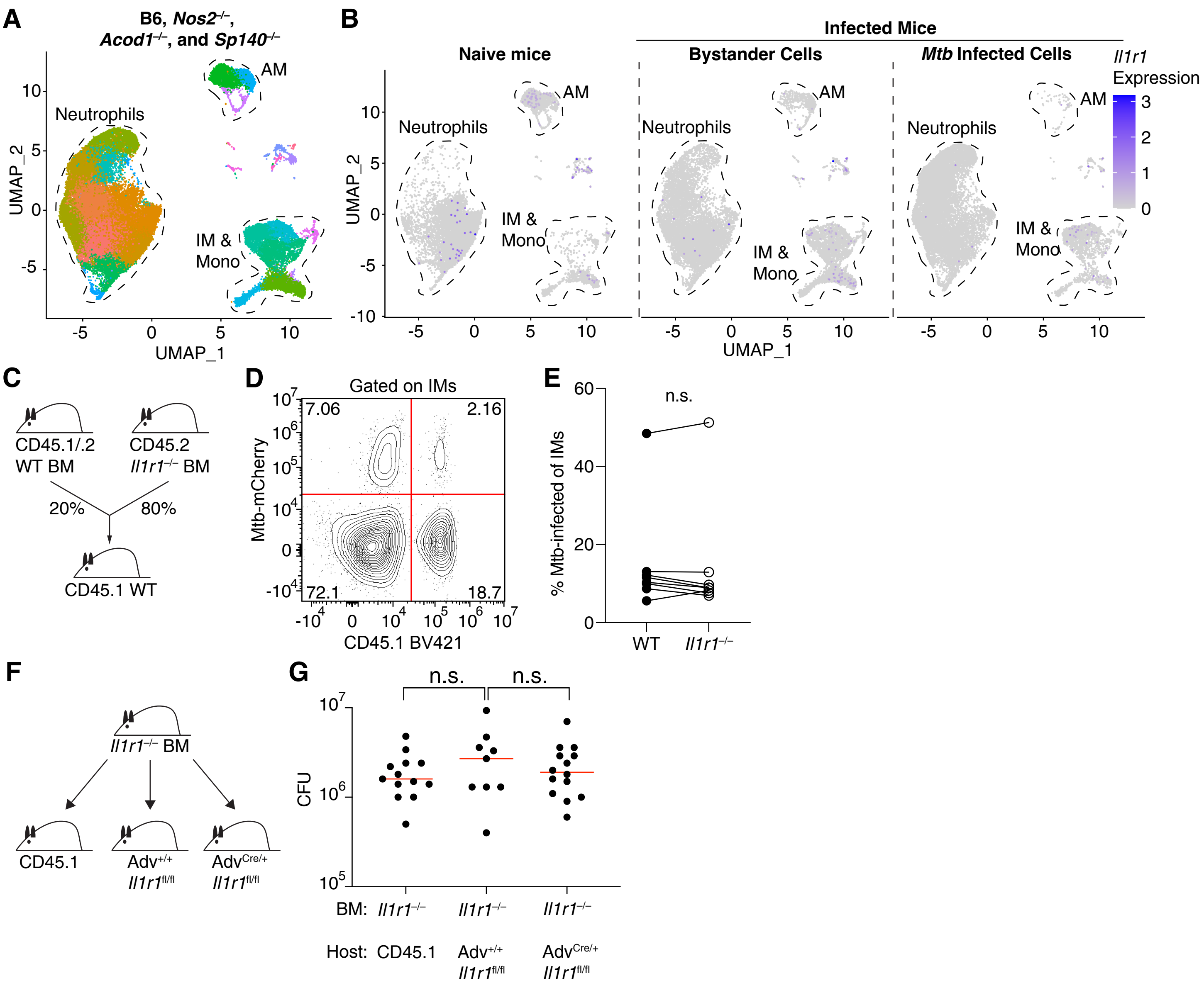


**Extended Data Figure 8.**

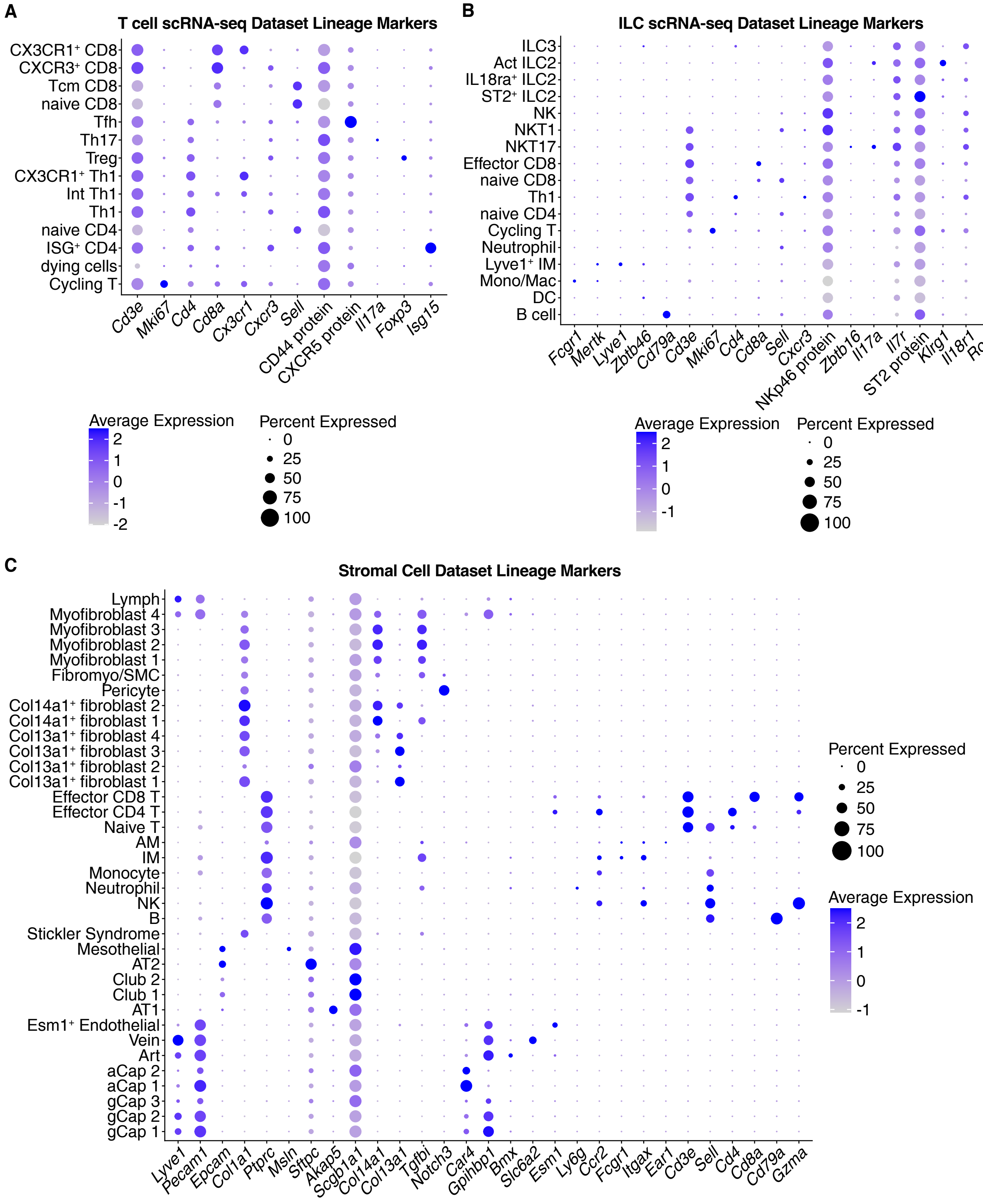


**Extended Data Figure 9.**

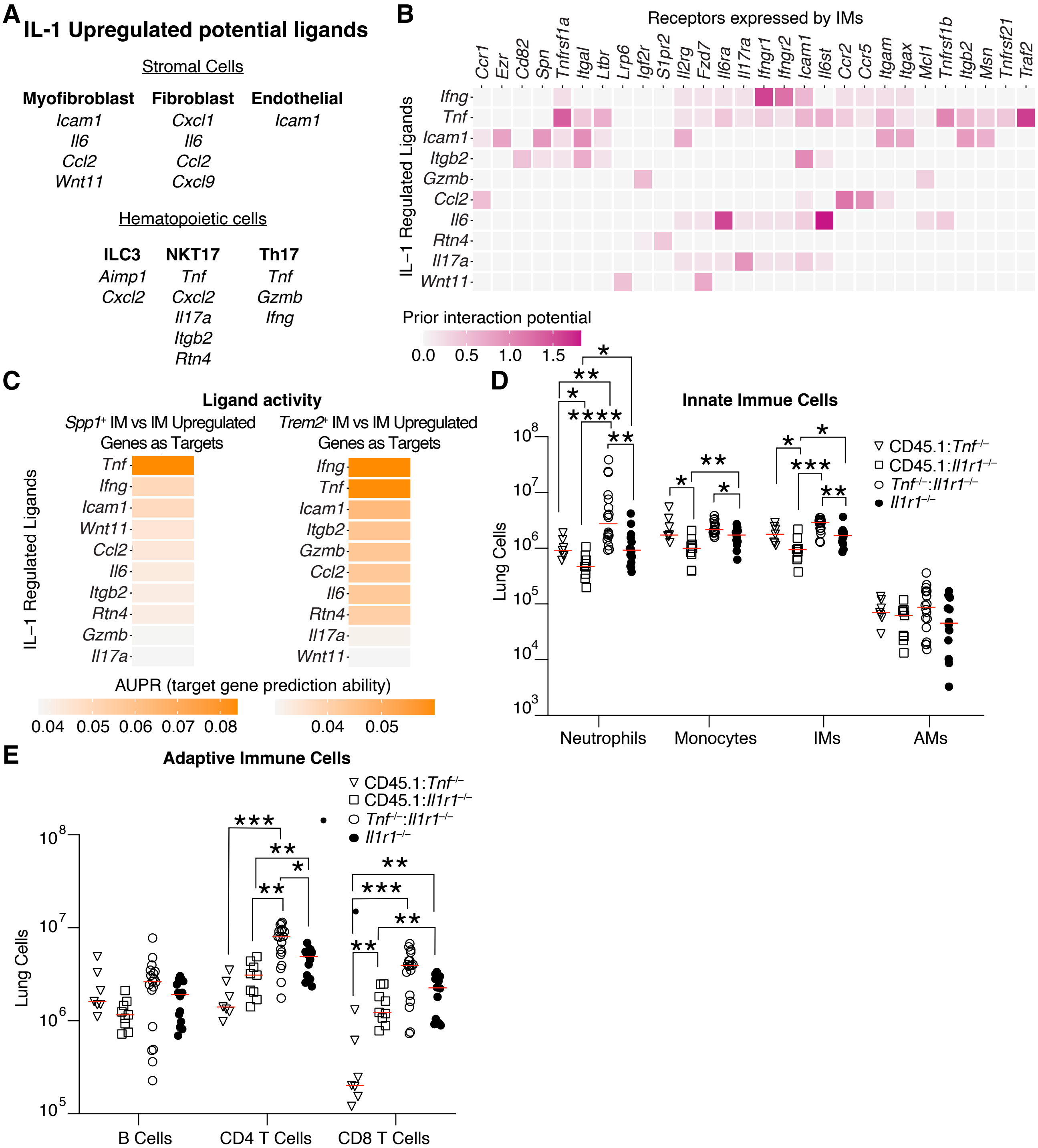


**Extended Data Figure 10.**
